# AI-guided cryo-EM probes a thermophilic cell-free system with succinyl-coA manufacturing capability

**DOI:** 10.1101/2022.10.08.511438

**Authors:** Ioannis Skalidis, Fotis L. Kyrilis, Christian Tüting, Farzad Hamdi, Toni K. Träger, Jaydeep Belapure, Gerd Hause, Marta Fratini, Francis J. O’Reilly, Ingo Heilmann, Juri Rappsilber, Panagiotis L. Kastritis

## Abstract

Cell-free systems display tremendous potential for biotechnological applications, complementing *in vitro* reconstituted enzymatic processes and traditional expression systems. However, they often represent “black boxes” without much insight into their components. Here, we characterize a thermophilic cell-free system that produces succinyl-CoA and discern its intrinsic, non-stochastic organization. By employing biochemical, biophysical, and bioinformatic methods we resolve its molecular composition, 3D architecture and molecular function at atomic resolution. We further report the high-resolution cryo-EM structure of the reaction’s main component, the oxoglutarate dehydrogenase complex core (E2o), which displays various structural adaptations. These include hydrogen bonding patterns confining interactions of participating enzymes (E1o-E2o-E3), electrostatic tunneling that drives inter-communication between subunits, and the presence of a flexible subunit, the E3BPo connecting E2o and E3. This multi-scale analysis of a cell-free system provides a blueprint for structure-function studies of complex mixtures of biotechnological value.

## Introduction

In-cell metabolic engineering is the go-to method for biotechnological production of various compounds of interest^1^. There are numerous successful examples^2, 3^, where, by engineering a specific biosynthetic pathway, the carbon flux to the desired product is increased, leading to strains with enhanced product generation capabilities^4^. Nevertheless, this methodology, even though widely applied, has clear disadvantages, some of them being: slow cell growth and wasteful biomass production^5^, an inherent environmental risk of genetic engineering^6^ or intolerance to the accumulation of toxic reaction products or by-products^7^. Cell-free system (CFS) technologies^8^ attempt to solve these issues by de-coupling product generation from the living organism, which bypasses synthetic limitations of the cell and its homeostatic mechanisms^9, 10^.

CFS technologies employ two main *in vitro* approaches. For the first, the desired biosynthetic pathway is reconstituted from individually purified components^11^, with the PURE system^12^ representing a highly successful example of this approach. Of course, this requires multiple expression, purification and fine-tuning strategies, each accompanied by its own obstacles^13^. The second approach is the use of crude cell lysates for product synthesis^14^. Here, substate is added to the lysate to begin generation of the desired product. This method, despite solving some of the issues mentioned above, also has significant requirements in *a priori* metabolic engineering of the desired microorganism strain to maximize production of the desired product and improve the stability of the desired enzymes^15^. An intriguing method that could greatly improve the second approach, besides genetic engineering, could be the use of fractionated native cell lysates^16^ derived from thermophilic organisms, which can be a potential source of stable and efficient enzymes due to their environmental adaptation. In parallel, size-exclusion chromatography provides a way of simplifying a native lysate and separates it into what can be subsequently seen as a “modular protein toolbox”: Each fraction will contain a distinct sub-population of the cell’s total protein content which can be characterized by a multitude of techniques^17, 18^ while being directly accessible to biochemical manipulation of in-extract biosynthetic pathways.

Succinic acid and L-lysine represent two important products with a wide range of biotechnological applications^19, 20^ and a constantly growing market demand, making their production an attractive target for cell-free methodologies. Both products can be formed from succinyl-coA through separate biosynthetic processes^21, 22^. Optimization of their production has so-far only focused on metabolic engineering of the yield in the respective organisms^23^ and not on *ex vivo* cell-free strategies. This is perhaps due to the deep interconnectivity with other primary metabolic pathways such as succinyl-coA production that require an active oxoglutarate dehydrogenase complex (OGDHc)^24^. OGDHc is one of the main regulators of metabolic flux in the Krebs cycle, meaning that cells maintain its abundance and activity under tight control^25, 26^. Furthermore, OGDHc is highly sensitive to reactive oxygen species (ROS)^27^, and its possible inhibition due to oxidative stress could prove detrimental to a cell’s overall metabolism. Clearly, OGDHc represents a bottleneck for metabolic engineering targeted towards increasing the yield of succinic acid, L-lysine, or other products, and is still not fully understood.

At the molecular level, OGDHc is a megadalton-size assembly of multiple copies of three enzymes, namely E1o (oxoglutarate dehydrogenase, EC 1.2.4.2), E2o (dihydrolipoyl succinyltransferase, EC 2.3.1.61) and E3 (dihydrolipoyl dehydrogenase, EC 1.8.1.4). E2o forms the core of the complex with a stable stoichiometry of 24 copies in a cubic assembly, whereas the E1o and E3 proteins assemble on its periphery to carry out the complete reaction in a yet stoichiometrically and structurally unknown high-order metabolon (**Fig. 1A**). Although OGDHc single enzyme components have been structurally resolved and kinetic characterization of its discrete steps are studied^28-30^ (**Fig. 1B**), their intercommunication is unknown, hampering, therefore, structure-function studies underlying this metabolon. This is due to the fact that the metabolon has not yet been reconstituted *in vitro* as a functional entity despite decades of research. In addition, the lipoyl domain (LD) which transfers lipoyl intermediates from one active site to the next one (E1o → E2o → E3) is part of a highly flexible E2o arm, representing a particular challenge for its study by traditional structural methods. The potential involvement of additional structural elements such as subunits that could contribute to OGDHc cross-communication, *i*.*e*., KGD4^31^, further complicate the complex’s characterization.

**Figure 1:**
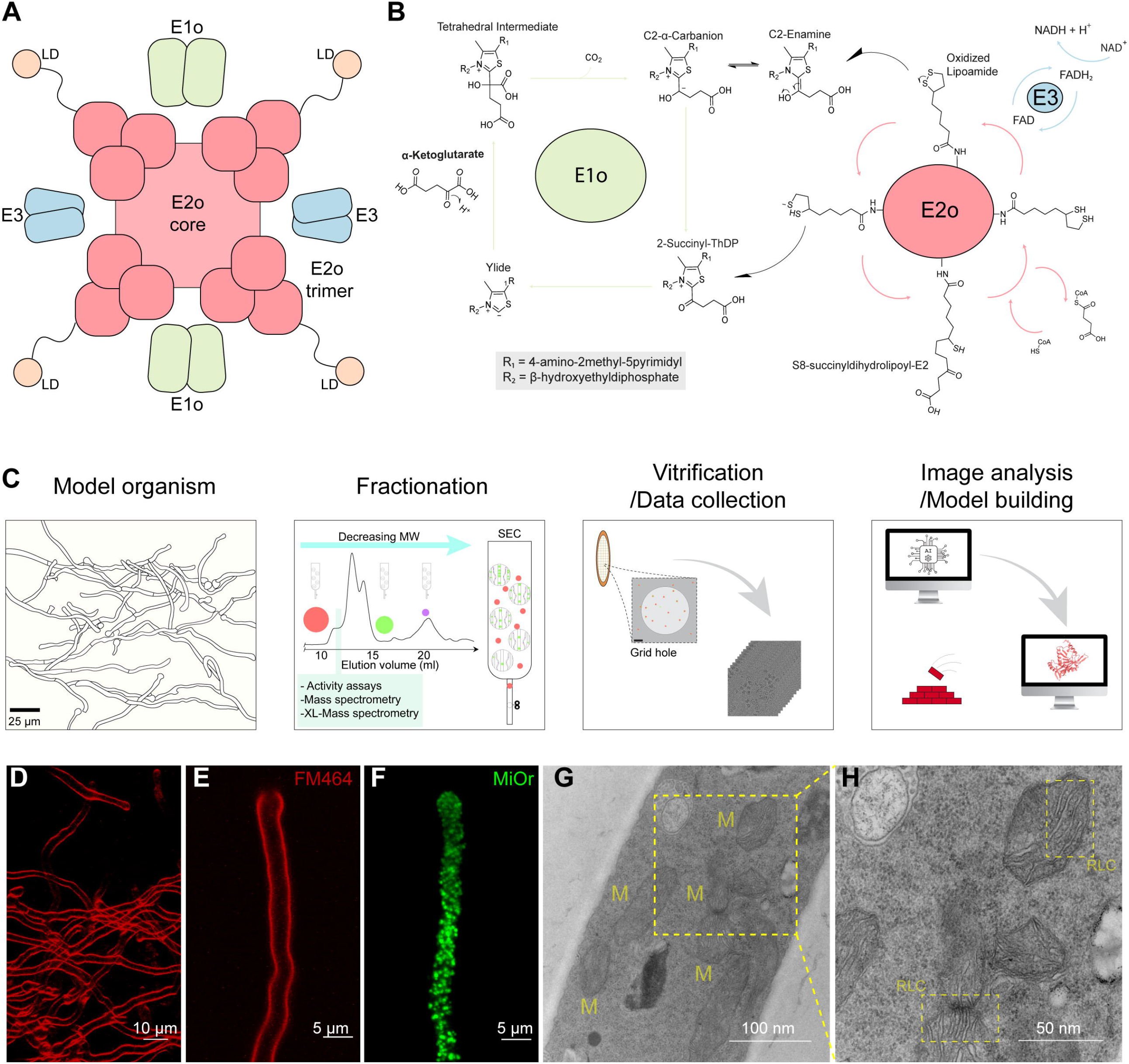
Previous knowledge on OGDHc, a workflow to expand it and the model organism that makes it possible. **(A)** Schematic representation of previous knowledge on OGDHc overall architecture. A cubic core comprised of 24 E2o proteins, with extended flexible linkers ending in the ordered LD domain, and E1o and E3 dimers on the periphery. **(B)** Complete reaction scheme of OGDHc. **(C)** Schematic representation of the workflow that is followed in order to produce the cell-free system and characterize it. **(D)** Fluorescent light microscopy of isolated *C. thermophilum* filaments, with the cell membrane in red color, stained with FM464. **(E)** Fluorescent light microscopy of a single *C. thermophilum* filament, with the cell membrane in red color, stained with FM464. **(F)** Fluorescent light microscopy of a single *C. thermophilum* filament, with the highly abundant mitochondrial content visible in green color, stained with MiOr. **(G)** Transmission electron microscopy image of cryo-fixated ultra-thin sections of *C. thermophilum* filament. With “M” the mitochondria are annotated in the image. **(H)** Zoom-in into the mitochondria ultrastructures visible in **(G)**. The ribbon-like christae of the *C. thermophilum* mitochondria are visible (annotated as “RLC”).

In this work, we endeavor to elucidate the structure of the OGDHc metabolon in the context of a cell-free native lysate fraction derived from *Chaetomium thermophilum* after a single-step fractionation of the crude lysate. *C. thermophilum* is a thermophilic, filamentous fungus with an optimal growth temperature of 54°C and the inherent thermostability of its proteins and protein complexes makes it an ideal model for structural studies focusing on multi-component targets, such as OGDHc or other protein complexes **(Fig. 1C)**. The inherent advantages of a thermophilic cell extract are put forth while we dissect its composition and architecture. We employ biochemical assays to reveal the complex’s kinetic parameters for producing succinyl-CoA, identify all its components and utilize cryo-electron microscopy, crosslinking and artificial intelligence (AI) to elucidate the mega-complex’s structure at high resolution with rigorous data collection strategies. Overall, we provide a multi-scale snapshot of a CFS that is underlined by flexible regions dictating transient enzyme-enzyme interactions, enzyme clusters that electrostatically channel OGDHc intermediates and by a higher-order architecture of OGDHc critical to further fathom the CFS’ function.

## Results

### Biochemical isolation of a cell-free system from a thermophilic, filamentous fungus

The thermophilic fungus *C. thermophilum* was grown at 54° C for 20 h before lysing and fractionating by size-exclusion chromatography (SEC) 8 *g* of cell fresh weight (**Methods**). *C. thermophilum* hyphae (**Fig. 1D-F**) are rich in mitochondria, as validated by confocal and electron microscopy methods (**Fig. 1F, G, H**). Mitochondria exhibit the expected lamellar, ribbon-like cristae, forming parallel stacks in cross section^32^ (**Fig. 1G, H**). A high abundance of mitochondria **(Fig. 1G,H)** is consistent with the increased respiratory rates of thermophiles^33^ and motivated us to derive a cell extract enriched in mitochondrial activities in which the functional OGDHc presents itself in high abundance^17^ with all its known subunits (E1o, E2o, E3)^34^. Here, we verified presence of all enzymes via WB **(Fig. 2A, Supplementary Fig. 1B)** and proceeded with quantitative kinetic parameter characterization for all of the metabolon’s soluble substrates, namely NAD+, α-ketoglutarate (α-KG) and coenzyme-A (CoA). OGDHc catalyzes the conversion of α-ketoglutarate (α-KG) to succinyl-CoA through a multiple-step reaction involving different co-factors (**Fig. 1B**): thiamine diphosphate (ThDP) binds to the E1o, a lipoate covalently attached to lipoyl-binding domain (LD) of the E2o, whereas FAD binds to its respective site at the E3. E1o and E2o are unique for this complex, but E3 is shared amongst all oxo-acid dehydrogenase complexes. In-fraction *K*_M_ values were determined at [149.80 ± 41.78] μM, [146.11 ± 46.15] μM and [22.81 ± 11.93] μM respectively (**Fig. 2B, Supplementary Fig. 1A, Supplementary Table 6**) and are comparable to those from other thermophilic counterparts^35, 36^. To accurately display the advantage of a CFS derived from a thermophilic eukaryote, we repeated the characterization and performed a comparison of the reaction velocity for α-KG in a temperature gradient between a *C. thermophilum* and a *S. cerevisiae* equivalent CFS (**Fig. 2B, Supplementary Table 6**). The derived data clearly displays an increase of reaction velocity for the *C. thermophilum*-derived CFS as contrasted by the velocity decrease of the yeast equivalent sample, showing the suitability of the CFS derived from a thermophilic organism for biotechnological application schemes that may also rely on high temperatures. Our findings demonstrate that a single fraction can be well-employed in CFS pipelines, is scalable due to its biochemical nature and is exploitable for product formation without the need for further purification or enrichment schemes.

**Figure 2:**
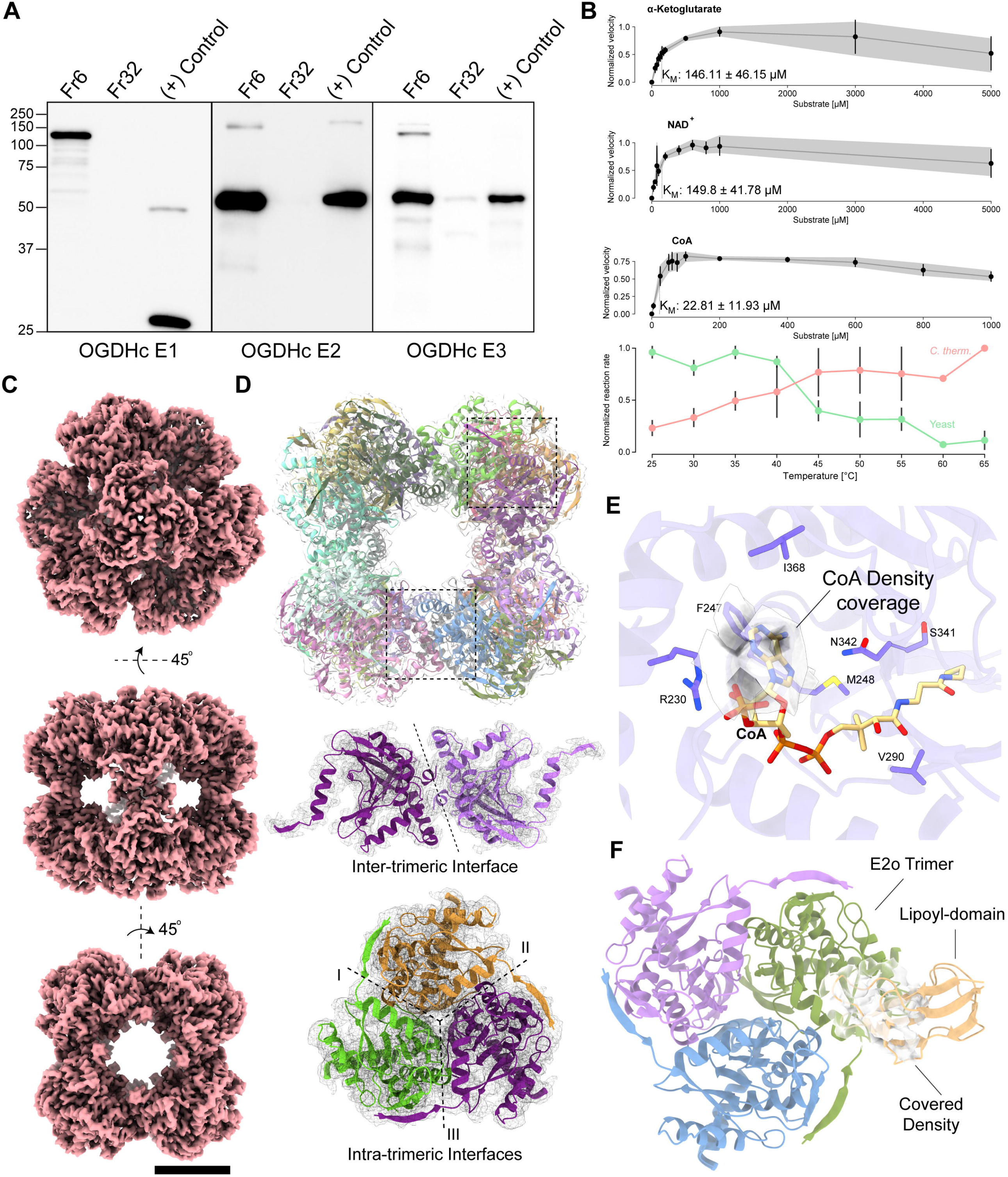
Biochemical characterization of OGDHc and the Cryo-EM structure of the OGDHc E2o 24-mer core. **(A)** Western blots displaying the detection of all OGDHc components in the cell-free system at the expected molecular weight (MW). A low-MW fraction was used as negative control, whereas the overexpressed protein that was used to create the antibodies was employed as a positive control. Due to the large size of the E1o protein, a fragment was employed for this reason (**Supplementary Table 7**). **(B)** Enzymatic characterization of the native cell-free system OGDHc. α-Ketoglutarate, NAD^+^ and CoA were used at the concentrations shown in each plot accordingly, the velocity was normalized to 0-1 and a line is connecting each point. The bottom graph displays the change in reaction velocity (after normalization to 0-1) in relation to temperature for *C. thermophilum* as compared to a yeast equivalent sample. The *K*_M_ values shown in the plot were obtained by the Burk-Lineweaver plots shown in **Supplementary Figure 1A** and the gray background for each graph and the black bars at the bottom graph represent the standard deviation derived from N=3 independent biological replicates and 2 technical duplicates for each replicate. All values shown here are listed in **Supplementary Table 6. (C)** The 3.35 Å (0.143 FSC) cryo-EM map of the *C. thermophilum* OGDHc E2o core. Scale bar: 5 nm. **(D)** The reconstructed model of the *C. thermophilum* E2o core, fitted in the cryo-EM map. Beneath, the inter- and intra-trimeric interfaces can be observed in isolation. **(E)** The CoA binding region of the E2o, with the annotated amino-acid sidechains that participate in its coordination. Additionally, resolved density in the pocket may correspond to a bound CoA. **(F)** Resolved density in the periphery of the E2o vertex trimer, possibly belonging to a bound E2o lipoyl domain.

### The 3.35 Å cryo-EM structure of the native, endogenous eukaryotic OGDHc core from the CFS elucidates structural features

To structurally characterize the OGDHc metabolon, we performed an intensive cryo-EM data acquisition of the characterized CFS. The structure determined for the E2o core reached 3.35 Å resolution (**Supplementary Fig. 2A, Supplementary Table 1**) enabling *de novo* model-building and considerably improving the resolution over that of the previously reported E2o core^18^ (**Supplementary Fig. 2D**). The EM map (**Fig. 2C**) displays higher-order structural features, *e*.*g*., inter-, and intra-trimeric interfaces (**Fig. 2D**), captures secondary structure elements and accurately places amino-acid side chains (**Supplementary Fig. 2C**). We were able to identify the E2o’s catalytic active site and fully model side-chain conformations of the amino-acids participating in CoA binding which are conserved across oxo-acid dehydrogenases. F247 has an outward facing orientation to accommodate the CoA **(Fig. 2E)**, interacting and stabilizing the CoA 3’, 5’-adenosine diphosphate group via π-stacking. The limited presence of electron density for the CoA pantoic acid, β-alanine, and cysteamine components underlies the inherent flexibility of endogenously bound coenzyme A with implications for function^37^.

Another density is observed proximal to the E2 lipoyl binding site **(Fig. 2F)**; in the bacterial endogenous pyruvate dehydrogenase complex (PDHc), which also forms a cubic core – a feature of keto-acid complexes across kingdoms – an equivalent density was attributed to the lipoyl domain (LD)^38^. Bound LD was also structurally refined in the eukaryotic PDHc E2 active site in a similar, stable conformation^39^. In the eukaryotic, native, and active OGDHc presented here, this density persists, albeit to lower resolution, and the LD can be partially accommodated **(Fig. 2F)**. A major difference lies in the fact that, compared to bacterial PDHc which could be trapped in a proposed resting state with all LDs bound to the E2 active sites, the present eukaryotic OGDHc highlights an alternative resting state where LDs are sub-stoichiometrically and transiently interacting with the E2 core. Overall, this OGDHc core structure comprises the highest resolution reconstruction reported to date for a protein community member characterized in native cell extracts. Besides obvious high-resolution features, the higher-order interfaces and LD/CoA active sites are defined at high resolution, while, due to its endogenous nature, low-resolution densities appear for the flexibly bound CoA and the sub-stoichiometrically-bound LD.

### Revealing E2o dihydrolipoyl succinyltransferase core compaction in the context of the native metabolon

The OGDHc dihydrolipoyl succinyltransferase (E2o) reconstruction highlights broad similarity with the human overexpressed, inactive counterpart in terms of overall fold but with localized adaptations (**Fig. 3**). The E2o *N-ter* in previously published structures does not acquire a specific fold (**Fig. 3A**). By contrast, in *C. thermophilum* this element obtains a β-strand (E188-M194) conformation which, along with a β-hairpin motif of residues N266-D282 creates an extended β-sheet (**Fig. 3B**). This adaptation possibly contributes to the stabilization of the core via extensive hydrogen bonding and affects subunit inter-communication in the higher-order state of the 24-meric cubic E2o core (**Fig. 6C**). This fold imposes a condensation of the E2o core trimer when compared to its previously published human counterpart, translated in a lower centroid distance across subunits (**Fig. 3C, Supplementary Fig. 3A**). Subsequent HADDOCK scoring of the human^29^ and C. *thermophilum* inter- and intra-trimeric interfaces (**Supplementary Fig. 3C, D, Methods**) show consistently higher scores for the native, thermophilic E2o interfaces. Considering that buried surface area is proportional to the experimentally measured dissociation constants (*K*_D_) of biomolecular complexes that do not undergo large conformational changes^40^, our results suggest a stronger binding of *C. thermophilum* E2o enzymes. In line with this notion, *C. thermophilum* intra- and inter-subunit interfaces are 1.7 and 1.5 times larger than the corresponding interfaces of the human E2o core (**Supplementary Fig. 3C, D**) and include extensive charge-charge interactions (**Supplementary Fig. 3C, D**). Another consequence of the afore-mentioned compaction is the induced confinement of the E2o *N-ter* LD. An antiparallel β-sheet has greater stability as compared to the previously resolved loop conformation^41^. Antiparallel and mixed sheets can withstand both distortions (twisting and β-bulges) and exposure to solvent; therefore, the newly identified structural element can serve as an anchor point for restraining conformations explored by the flexible region connecting the two ordered E2o domains, *i*.*e*., the succinyltransferase forming the E2 core and the LDs carrying the intermediates.

**Figure 3:**
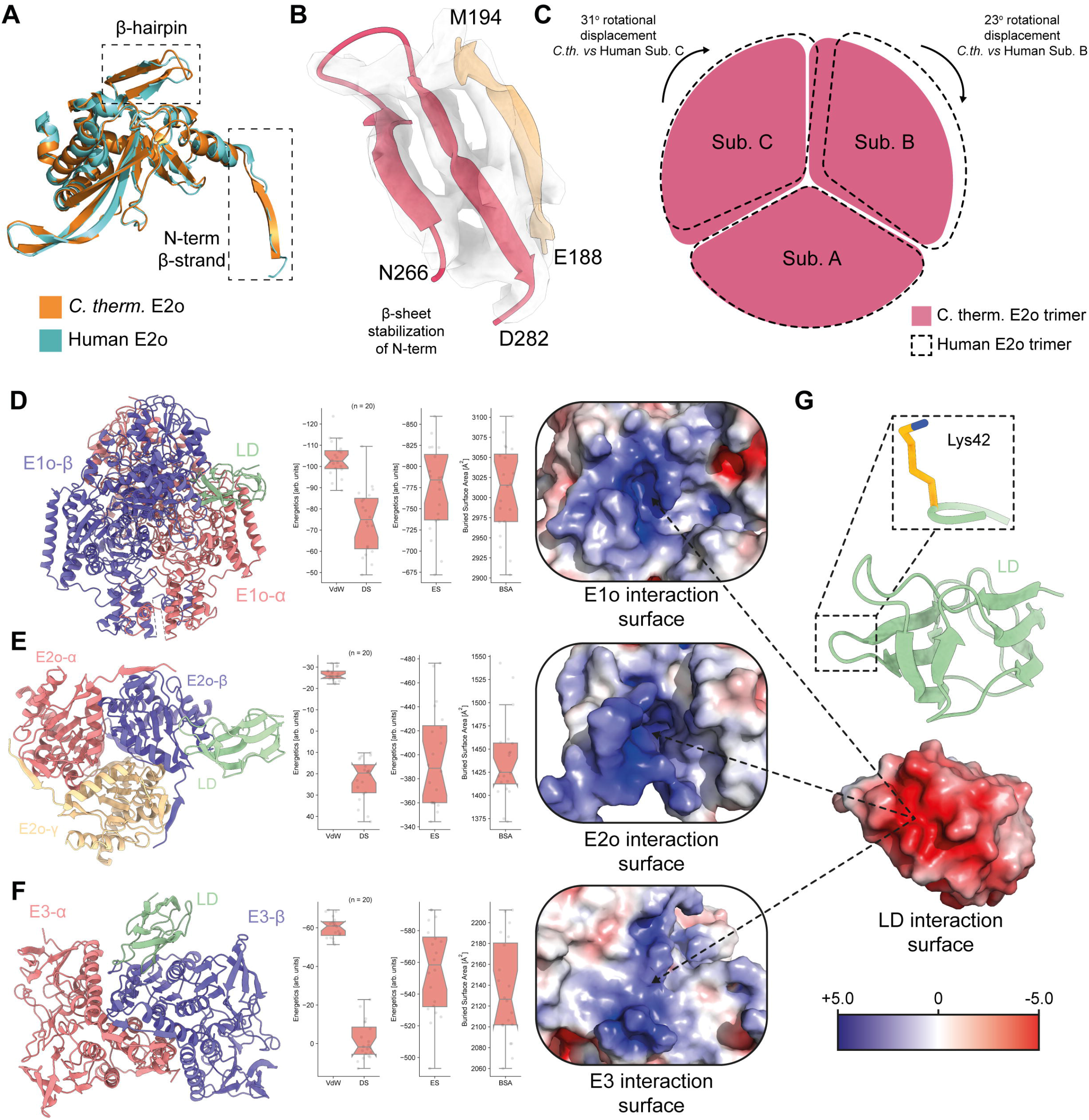
Structural adaptations of the *C. thermophilum* E2o in comparison to its mesophilic counterpart and electrostatic interaction amongst the OGDHc components. **(A)** A *C. thermophilum* E2o protein (orange) aligned with its human counterpart (blue). Noticeable differences include a tighter turn conformation resulting in better alignment of the *C. thermophilum* β-hairpin model and the β-strand conformation of the *C. thermophilum N-ter*. **(B)** The β-hairpin motif of N266-D282, along with the *N-ter* β-strand E188-M194 form a β-sheet that contributes to the core’s overall stability. **(C)** Schematic representation of the human E2o vertex trimer rotational displacement in comparison to the experimentally resolved *C. thermophilum* E2o trimer, displaying a “loose” conformation for the mesophilic counterpart. **(D)** AI-derived model of the E1o-LD interaction. Overall model, energetics (VdW=Van der Waals interactions, DS=Desolvation energy, ES= Electrostatics) and buried surface area (BSA), and the overall charge of its interaction interface with the LD can be seen in left, middle and right respectively. **(E)** AI-derived model of the E2o-LD interaction. Overall model, energetics and buried surface area, and the overall charge of its interaction interface with the LD can be seen in left, middle and right respectively. **(F)** AI-derived model of the E3-LD interaction. Overall model, energetics and buried surface area, and the overall charge of its interaction interface with the LD can be seen in left, middle and right respectively. **(G)** AI-derived model of the ordered LD domain of E2o. Lys42 carries the lipoyl-moiety. A highly negatively charged interaction interface is observable.

### Electrostatic complementarity of comparable magnitude governs the 3D architecture of the OGDHc reaction interfaces

During catalysis (**Fig. 1B**) in the active OGDHc metabolon that we captured (**Fig. 2B**), the flexible E2o arm holding the LD shuttles the succinyl-intermediate from the E1o to the E2o active sites, and is re-oxidized by the E3^42^; Therefore, the lipoylated lysine of the LD must have unhindered access to each enzymatic site. To structurally characterize these transient interfaces, AlphaFold2-Multimer modelling of the three metabolon-embedded component interfaces (E1o-LD, E2o-LD and E3-LD) was performed. The models generated for all three complexes **(Fig. 3D-F, Methods)** are of high quality, and their predicted interfaces confidently rank within experimental error (**Supplementary Fig. 5A, B**). In addition, (a) the fit of the LD in the proximal density of the E2o core derived by cryo-EM (**Fig. 2F**) is similar to the one predicted by AI in terms of positioning (**Fig. 3E**) and (b) the distance of the lipoylated lysine to the cofactor or active site for all generated interfaces is in the range of 10 to 25 Å (**Supplementary Fig. 6A-D**). This range corresponds to the length of the lysine side chain and the lipoate and also reflects the intrinsic flexibility of the interacting proteins^43^, also observed in a recent bacterial PDHc E2o-LD interface^39^. E1o forms a homodimer (**Fig. 3D**), with two thiamine diphosphates (ThDPs), and potentially two LD binding sites localized at the E1o inter-chain interface (**Fig. 3D**). The lipoylated lysine distance to the ThDP C2 atom that is succinylated during α-ketoglutarate decarboxylation is biochemically feasible (*d*=14 Å) (**Supplementary Fig. 6A**). E2o binds a single LD per monomer (**Fig. 3E**) and the lipoyl-lysine is proximal to the bound CoA, with a lysine Cα-CoA thiol group distance of *d*=14 Å as well (**Supplementary Fig. 6B**). Lastly, the E3 active site locates a conserved disulfide bridge and a histidine in proximity involved in the E3-mediated LD-re-oxidation reaction^44-46^. The AI-based E3-LD complex shows that the lipoylated lysine is in reasonable distance to access disulfide bonding in the E3 active site (*d*=23 Å) (**Supplementary Fig. 5D**). Note that a second, alternate conformation model for both E1o-LD and E3-LD complexes was generated (**Supplementary Fig. 5C, D**) but did not satisfy the above-mentioned biochemical constraints essential for an active metabolon (**Fig. 2B**).

The biochemically-(E1o-LD, E2o-LD and E3-LD) and cryo-EM-(E2o-LD) validated AI-generated metabolon-embedded interfaces share a common characteristic, revealed after subjecting each to energy-based refinement (**Fig. 3D-F**): All interfaces are governed by strong electrostatic interactions of comparable magnitude (**Fig. 3D-F**) that comprise a signature energetic component for transient interfaces with high on/off rates^47^. Complementary electrostatics (**Fig. 3G, Supplementary Fig. 3B**) are a collective characteristic in metabolons shuttling intermediates via a “swinging arm” mechanism^48^ and other metabolic processes requiring rapid on/off switching^49, 50^. Indeed, electrostatic potential maps highlight extensive positively charged regions for the LD binders (**Fig. 3G**). These surfaces attract and accommodate the LD’s negatively charged surface and overall contribute to the structural compaction of oxo acid dehydrogenase complexes^51^ that is a key principle to achieve metabolic channeling via the “swinging arm” mechanism^52^.

### Mapping the protein communities observed in a CFS and identification of OGDHc E3BP

To elucidate the broader proteome and its interactions within the CFS, we performed mass spectrometry-based crosslinking experiments directly in the extract (**Methods**) while benchmarking crosslinker concentrations (**Supplementary Fig. 1C**). Overall, with an FDR of 2% at the peptide recovery level **(Methods)**, we retrieved 4949 residue-residue crosslinks (3 632 intra- and 1 317 inter-residue crosslinks), of which 99.4% mapped to *C. thermophilum* polypeptide chains (**Supplementary Table 2**). Out of a total of 2091 polypeptide chains, 505 polypeptide chains were crosslinked across 2 biological and 2 technical replicates (**Supplementary Table 2**). This shows that 29.1% of the total *C. thermophilum* proteome organizes in cellular communities^53^ which are significantly larger than previously described^34^.

Network analysis identified 54 cellular communities from diverse cellular compartments (**Supplementary Table 2**) in which 1488 identified members participate. Subunit coverage across communities is high (67%+/-24%) and includes examples such as the complete cytoplasmic (N=128, 96% of subunits) and mitochondrial (N=74, 99% of subunits) translation apparatus. Additionally, we identify all subunits of the pyruvate dehydrogenase/tricarboxylic acid cycle community crosslinked together (PDHc/TCA, N=25, 100% of subunits, **Supplementary Table 2**). Within the PDHc/TCA community, crosslinks define interactions within and across eleven participating enzymes (122 intra- and 169 inter-links, **Supplementary Table 2**). The E3 protein displayed the largest number of inter-crosslinks identified in the experiment (N=88) and was found in proximity to seven other community members (**Fig. 4A**), indicating its diverse role in mitochondrial metabolism; E3 proteins explore proximal spatial positions with all subunits of PDHc (E1p, E2p, E3BP), OGDHc (E1o, E2o), and a subunit of the hypothesized mitochondrial heme metabolon^54^, interacting within the latter with a putative holocytochrome c synthase (HCC) **(Fig. 4A)**. However, sequence analysis highlighted a fused, misannotated ribosomal protein at the HCC *N-ter* (**Supplementary Fig. 5C**) that corresponds to the elusive eukaryotic KGD4 OGHDc subunit^31^ that tethers the E3 to the OGDHc E2 core (residues M1-T130). This *N-ter* shares high similarity with KGD4 from other eukaryotes (**Supplementary Fig. 5C**). Reanalysis of MS data^34^ across cellular fractions shows strong co-elution of KGD4 with the rest of the OGDHc subunits in all three biological replicates (**Supplementary Table 2**). Because its function is essentially the same as the PDHc E3BP -tethering E3 to the E2 core-we designated the co-eluting protein as the *C. thermophilum* protein E3BPo.

**Figure 4:**
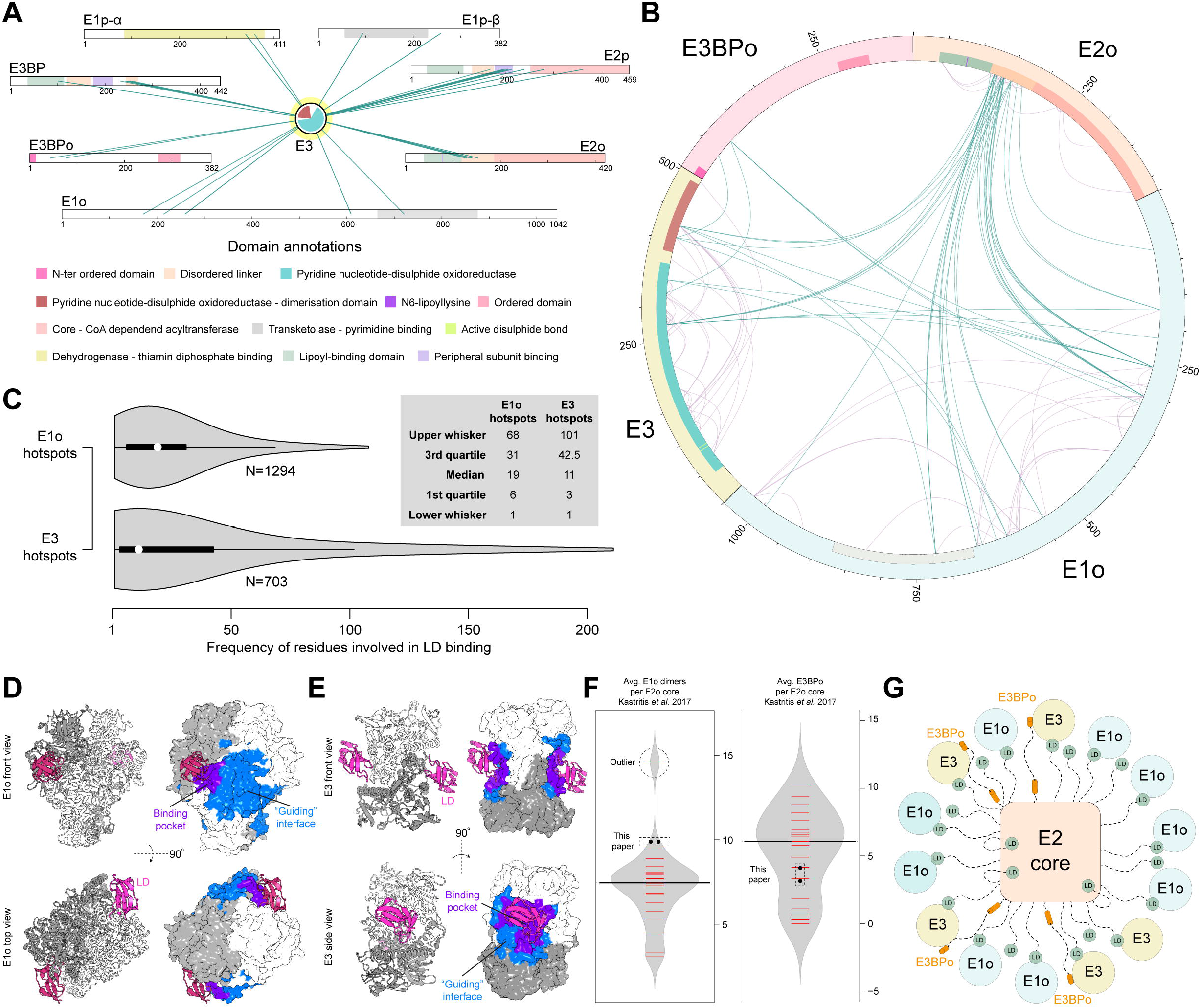
Crosslinking mass spectrometry (crosslinking-MS), crosslinking MS driven docking and MS stoichiometric calculations of OGDHc protein components. **(A)** Inter-crosslinks of E3 reveal a plethora of interaction partners. **(B)** The OGDHc protein components are highly interconnected. Blue lines represent the inter-protein crosslinks and purple the intra-protein crosslinks. **(C)** Frequency plots of residues involved in the binding interface between the LD and E1o (upper) and LD and E3 (lower). **(D)** Residue frequencies of each residue involved in the interaction between the LD (pink) and the E1o (gray) reveal a “guiding” interface (blue) between the two that orients the LD towards its binding pocket (purple). **(E)** Residue frequencies of each residue involved in the interaction between the LD (pink) and the E3 (gray) reveal a “guiding” interface (blue) between the two that orients the LD towards its binding pocket (purple). **(F)** Stoichiometric calculations of previously published MS data^34^ and newly derived MS data reveal the stoichiometry of the E1o and E3BPo components that take part in the formation of the fully active, native OGDHc. **(G)** Schematic representation based on the stoichiometries derived from the MS data presented in this article. The 24-mer E2o core is surrounded by at least 4 E3 dimers that are bound to 4 E3BPo, joined by 9 E1o dimers.

### A crosslinking informed network elucidates interactions within the OGDHc metabolon

44 inter- and 81 intra-molecular unique crosslinks were identified in the OGDHc components (E1o, E2o, E3, E3BPo) (**Fig. 4B**). OGDHc is relatively compact in the cell free system, and enzymes not known to form physical interfaces (E1o/E3; E3BPo/E1o) are still present in relative proximity, as evident from their identified inter-crosslinks **(Fig. 4B)**. The relative proximity of E1o and E3 to the E2o core point to the involvement of the *N-ter* flexible region of the E2 localizing downstream from the LD in the primary sequence (**Fig. 4B**). This result led us to utilize crosslinking-driven flexible molecular docking of the previously validated interaction models of the LD with the E1o and the E3 to elucidate interaction surfaces sampled by the LD (**Fig. 4C-E**).

We calculated the frequency of E1o or E3 residues involved in LD binding from the docking results after adapting our docking protocols for incorporating crosslinks mapping to disordered regions proximal to ordered domains in protein-protein interactions (**Methods**,^40^). Mapping the top 25% of residues involved in LD binding derived from 400 explicit-solvent refined molecular models illustrates a distinct attraction surface for the LD to eventually bind to the respective active sites (**Fig. 4D, E**). Active site residues are identified as frequently contacted for both E1o (**Fig. 4D**) and E3 (**Fig. 4E**), but the attracting surface for the E1o is more diffuse in terms of recovered hot-spots as compared to the one calculated for the E3 (**Fig. 4D, E**). This higher signal diffusion for LD binding in the E1o suggests that the proximal surface may guide the LD towards the E1o binding pocket as it is rather deeply buried (**Fig. 4D**). For the E3, instead, the “guiding surface” is much more confined (**Fig. 4E**) at a narrow space equilaterally extending ∼5 Å from the flat E3 active site. These results point to distinct LD attraction mechanisms for the peripheral E1o and E3 subunits governed by flexible regions positioned downstream from the LD.

Intra-crosslinks identified not only validate the presence of the participating proteins, but also confirm that they form multimeric structures; intra-crosslinks for E3BPo were not identified and we hypothesize that it must exist as a monomer; per E3BPo copy, a single E3 is tethered via its flexible *N-ter* region (**Fig. 4B**) to the E2 core^31^. Translating iBAQ scores into qualitative stoichiometric data as previously done for PDHc^17^ points to an exact stoichiometry of 24 E2o subunits, validated by cryo-EM; and relative stoichiometries of <10 E1o dimers, and ∼4 E3BPo, tethering 4 E3 dimers in total (**Fig. 4F, G**). Because E3 is also in proximity to the E2o disordered regions and its proximal E2o LD domain, it is well possible that more E3s can be accommodated. However, considering that the E2 core may recruit up to 48 molecules, totaling a metabolon of 96 polypeptide chains (24 E2o and 24 E3BPo monomers, each tethering 12 E1o and 12 E3 dimers, respectively), relative stoichiometries reveal substoichiometric composition of the peripheral OGDHc subunits. This points to the presence of unbound flexible *N-ter* E2o core regions involved in LD trafficking.

### Cryo-EM and computational analysis of E1o, E2 and E3 proximity in the context of the metabolon

To elucidate the higher-order architecture of the OGDHc within the CFS, we performed asymmetric reconstructions from the single-particle data, leading to 2D projections where external densities were visible **(Fig. 5A)**. In these external densities, the identified E1o and E3 dimers should be located, each tethered via either the flexible regions of the *N-ter* of the E2o (**Fig. 4B**) or, in the case of E3, the flexible E3BPo (**Fig. 4B**, ^31^). The cryo-EM map (**Fig. 5B**) was determined at 21 Å resolution (**Supplementary Fig. 2B, Supplementary Table 1**), exhibiting densities around the E2o core, in a clustered and not in a diffused fashion **(Fig. 5B)**. Increasing density thresholds of the cryo-EM map showed additional, weaker densities (**Fig. 5B**). Systematic fitting of AlphaFold2-derived E1o-LD and E3-LD models of *C. thermophilum* (**Fig. 5B, Supplementary Figure 4A, B**) in the cryo-EM map resolved those molecules’ localization and suggests they are present in the periphery and not on or within the core densities (**Supplementary Table 3**). In addition, persistent docking solutions that recapitulated the positions of E1o and E3 in the map were calculated from the systematically fitted structural models associated with high cross-correlation values (**Supplementary Table 3, Fig. 5B**): Overall, 2 E1o and 3 E3 dimers could be resolved within the calculated external densities.

**Figure 5:**
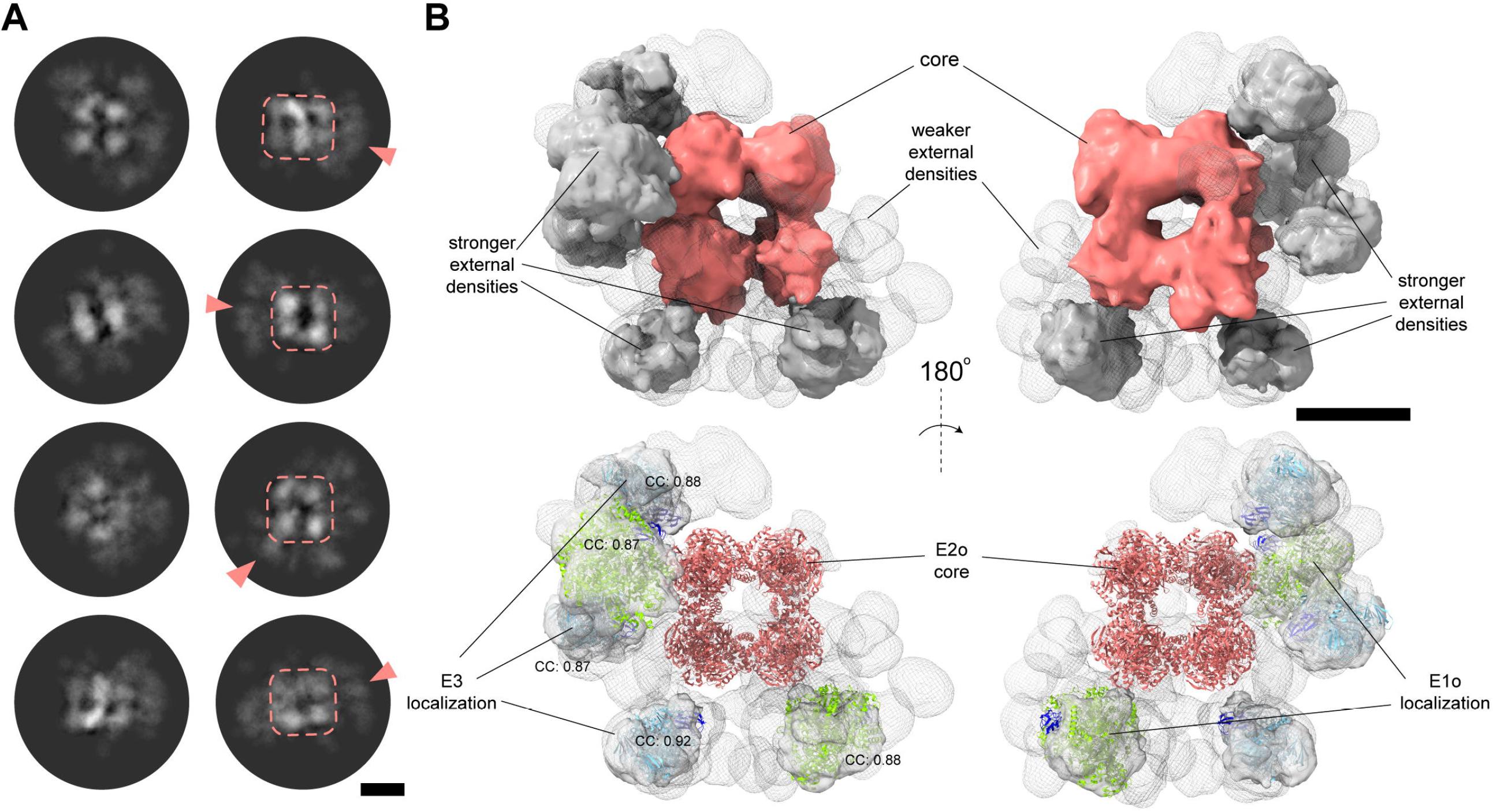
The asymmetric reconstruction of the complete OGDHc allows for the localization of its external subunits. **(A)** 2D class averages of the OGDHc present a core of high signal, surrounded by more diffused but still well-defined signal on the periphery. Scale bar: 5 nm. **(B)** The asymmetric 3D reconstruction of the OGDHc displays a strong density corresponding to the E2o core, and a mix of weaker and stronger external densities. In the stronger densities, E1o and E3 dimers can be localized with high confidence. Scale bar: 10 nm.

While the tethered E1o and E3 enzymes are asymmetrically distributed in proximity to the E2o core, they retain approximately equal distances from the core itself (**Fig. 5B**). The inherent flexibility of the tethering region in OGDHc does not reconcile with physical-chemical or statistical calculations for flexible protein regions of various categories (*e*.*g*., IDPs, molten globule, fully extended; **Supplementary Fig. 4C**). According to the E2o linker length measured in our derived 3D reconstruction **(Fig. 6A, Supplementary Fig. 4C)**, OGDHc flexible regions have properties in-between calculated lengths of “stretched” IDPs (IDPs as defined in Marsh and Forman-Kay^55^) and “constricted” linkers (as defined by George and Heringa^56^). To elucidate whether distances measured between ordered OGDHc participating enzymes that are mediated by flexible regions can be recapitulated in the available structural data, we calculated the distance of amino acid residues that are resolved but are sequence-wise further apart due to an unresolved stretch in the structural data. While various technical reasons may account for the presence of this unresolved stretch, it may also be an indicator of structural disorder^57^. By analyzing all Protein Data Bank (PDB)^58^ structural data as of June 2022 (191 144 structures), results show dramatically confined distances between the ordered residues preceding and following the “missing” stretch (**Fig. 6A**). This means that current structural data in the PDB very rarely include interactions of the kind reported in our results **(Fig. 5B, Fig. 6A)**. A relative “flattening” of calculated distances for absent polypeptide stretches of more than 25 amino acid residues is evident (**Fig. 5A**), and maximum Ca-Ca distances reported average at ∼30 Å, which are substantially shorter than those in the OGDHc metabolon.

**Figure 6:**
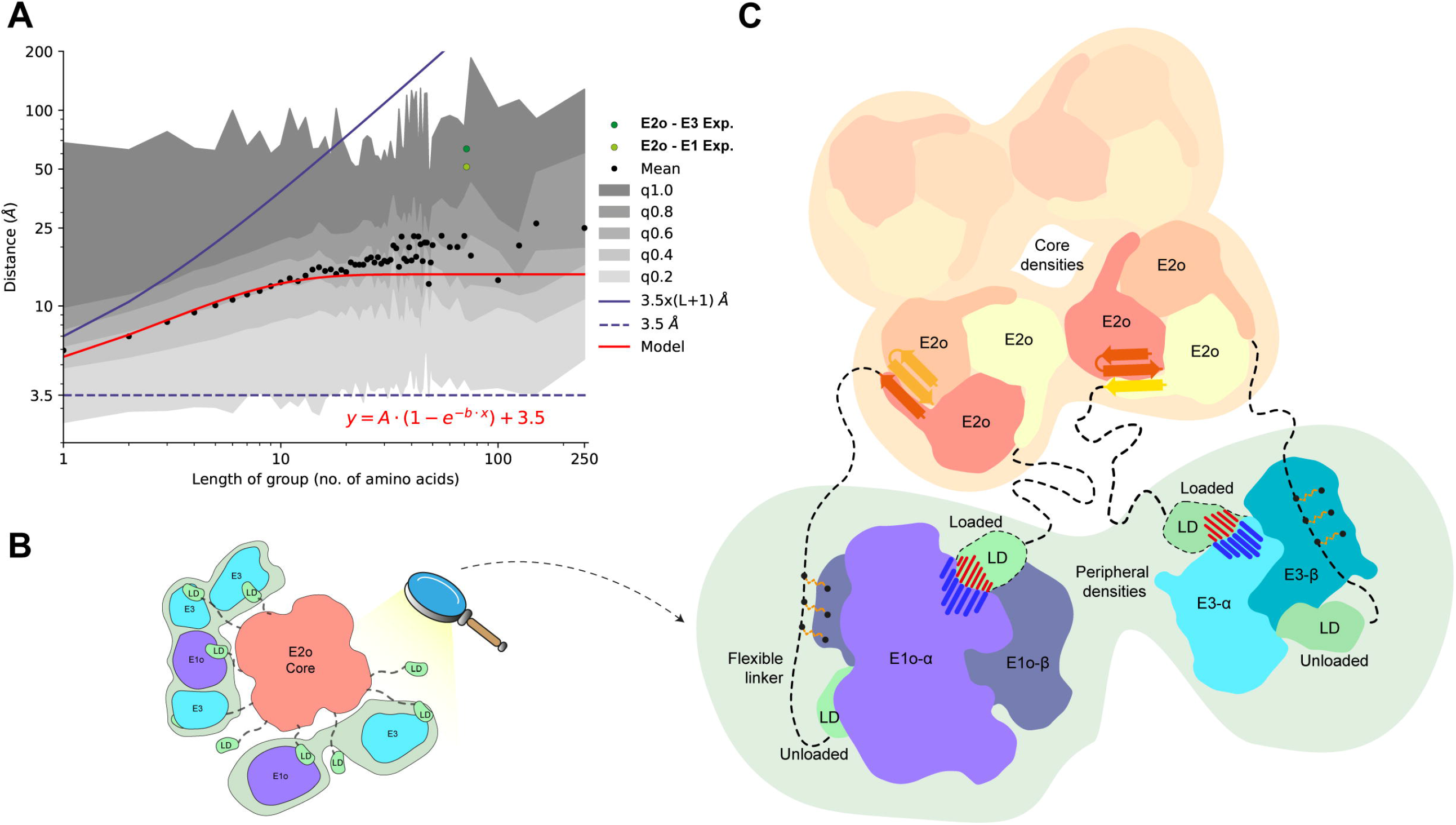
Flexible linker distances and a comprehensive model for the organization of the OGDHc. **(A)** Graph representing the mean distance values of the binned groups of all unresolved amino-acid sequences belonging to all protein structures deposited in the PDB. Black dots represent the mean value of a length of amino-acids group, with varying scales of gray the standard deviation after the integration of each quartile of total data, as denoted in the plot legend. The red line represents a fitted model that describes the relationship between the distance and the length of amino-acid groups. With light green the experimentally measured average distance between the *N-ter* of the resolved core domain of the E2o and the *C-ter* of the LD that is bound to a fitted E1o dimer on the periphery, while dark green represents the same average distance experimentally measured between the *N-ter* of the resolved core domain of the E2o and the *C-ter* of the LD that is bound to a fitted E3 dimer. The blue line represents the theoretical distance of an amino-acid sequence, while the dashed blue line represents the theoretical lower limit of any amino-acid sequence. **(B)** Schematic representation of the peripheral subunit organization of the resolved OGDHc. **(C)** Model describing all the novel observations concerning the organization of the OGDHc. The *N-ter* β-sheet conformation of the E2o core vertex trimer helps stabilize and compact the 24-mer E2o core, while simultaneously orienting the flexible linker that connects the LD domain. Cross-linking data reveals a fairly stable interaction between the flexible linker and the peripheral subunits, possibly hinting at a structural role of an un-loaded LD domain, while another, loaded LD domain performs the reaction cycling. The interaction between the LD and the peripheral subunits is governed by strong electrostatic forces.

## Discussion

Native cell lysate fractions provide a previously unexplored framework for the identification, characterization and optimization of enzymatic reactions in the context of cell-free biotechnology. Previously, their utilization has mostly been confined to the production of specific products of biotechnological relevance^8^, and has required metabolic engineering of the desired production pathway^21^ in order to maximize the yield obtained in specific organisms. Additionally, the majority of the characterizations of the enzymes involved in any metabolic pathway is usually performed *in vitro*^59^, thus omitting the enzyme’s “native” context. In consequence, identification, characterization and structural interrogation of the derived extracts is often lacking, hindering the understanding of the function within a complex assembly and utilizing these systems as “black boxes”. Our integrative results obtained by a broad range of analytical methods prove that CFSs can be interrogated with multiple approaches to eventually better understand their biochemistry. Importantly, a single-step fractionated cell lysate retains the “native” principles necessary to fully dissect a reaction pathway, both biochemically and structurally, making it possible to directly employ it in biotechnological applications.

Our combined findings expand our understanding of the proposed architecture of the oxoglutarate dehydrogenase complex in the context of a cell-free succinyl-CoA production system **(Fig. 6B, C)**. The complex’s core is a robust cubic ultrastructure where the E2o trimers that comprise the vertices are densely packed with the help of multi-subunit secondary structure elements (core domain *N-ter* β-sheet). The same structural element confines the available space of the extended *N-ter* flexible linkers that tether the LD domain which is the crucial domain for the transport of the substrates across sequential active sites that take part in the reaction, at around 50-100 Å for the LD-E1o interaction and 60-70 Å for the LD-E3 interaction. The flexible linker and LD of E2o, based on our XL-MS data, may also act as an “anchor” that, in the absence of specific peripheral subunit binding domains in the E2o *N-ter* sequence, maintains the E1o and E3 subunits in proximity to the E2o core. The disordered *N-ter* domain of E1o was recently resolved to interact with the E1o-LD binding site^60^ and may further aid in organizing E1o in proximity to the E2o core by interacting with other LD-binding interfaces in the metabolon. Both E1o and E3, being homodimers, contain dual LD-binding sites suggesting one playing an active role in the reaction, while the other can maintain a highly-charged complementary electrostatic tethering with a proximal “unloaded” LD domain. This “unloaded” LD domain may participate as an “anchor”, possibly also assisted by other weaker interactions of the flexible linker with surface residues of the peripheral subunits as well as the identified E3BPo protein, also captured in our XL-MS data to organize the E3 protein. The “structural” role of the extra LD domains can also be hinted by the sub-stoichiometric relationship between the core E2o and the peripheral E1o-E3-E3BPo subunits as quantified by the MS data presented here.

Overall, in this work, we characterized the complete contents of a CFS with succinyl-CoA producing capabilities, and elucidated the biochemical function and novel structural features of the reaction’s main player, the OGDHc and its components, E1o, E2o, E3, E3BPo. Our work gives rise to questions that need to be addressed: Is there in-fraction interplay among the multiple protein communities present? How can their biochemical and structural relationships be characterized? These questions can only be answered by employing automated, high-throughput cryo-EM of a fully, high-density, crosslinked CFS, combined with more large-scale data collection to tackle the issues of complex flexibility and relative low abundance of individual elements. Advances in instrumentation^61^, data collection^62^ and analysis^63, 64^, as well as the advent of precise AI-driven modelling^65^, will allow for the streamlining of the characterization of CFSs derived from native cell extracts and it is highly possible that in the near future our approach will lead to new paradigms of enzymatic analysis, optimization and biotechnological production of compounds to higher yields.

## Materials and Methods

### Model organism culture

*Chaetomium thermophilum var. thermophilum* La Touche 1950 (*Thermochaetoides thermophila*^66^) was acquired from DSMZ (Leibniz Institute DSMZ-German Collection of Microorganisms and Cell Cultures, Germany), and the preserved in freeze-dried ampoule spores were cultivated as directed by company guidelines (DSMZ Media list Medium 188, temperature: 45°C). After initial cultivation, the mycelium was propagated in liquid Complete Culture Media (CCM), containing per 1000 mL of ddH_2_O: 5.00 g tryptone, 1.00 g peptone, 1.00 g yeast extract, 15.00 g dextrin, 3.00 g sucrose, 0.50 g MgSO_4_ x 7 H_2_O, 0.50 g NaCl, 0.65 g K_2_HPO_4_ x 3 H_2_O and 0.01 g Fe_2_(SO_4_)^3^ x H_2_O. CCM final pH was adjusted to 7.1. Final cultures were performed as follows: Solid media plate cultures: liquid CCM was supplemented with 15.00 g of Agar/1000 mL ddH_2_O and the plates were then inoculated with mycelium and grown at 54°C. Liquid media cultures: 2000 mL Erlenmeyer flasks were filled to 40% of total volume (800 mL) with liquid CCM media, small pieces of freshly grown mycelium from Agar plates were added and then incubated under shaking at 110 rpm and 10% CO_2_ for 20 hours.

### Cell imaging

Fluorescence microscopy imaging was performed as follows: A Zeiss LSM880 (Carl Zeiss, Germany) with a 40X objective lens without immersion (plan-Apochromat 40X/0.95 N.A.) was used to image the samples and images were acquired with the ZEN Black image analysis software (Carl Zeiss, Germany). For membrane staining with FM 4-64 (ThermoFisher Scientific, USA), the dye was diluted in DMSO to a stock concentration of 10 mM; 1μl of stock solution was then added to 1 ml of *C. thermophilum* liquid cell culture, reaching a working concentration of 10 μM. Samples were imaged after 2 to 3 m of incubation time. For mitochondria staining with MitoTracker orange (ThermoFisher Scientific, USA) the dye was diluted in DMSO to a stock concentration of 10 μM; 5 μl of stock solution were then added to 1 ml of *C. thermophilum* liquid cell culture, reaching a working concentration of 50 nM. Samples were imaged after 10 m of incubation time. Transmission electron microscopy (TEM) imaging was performed as follows: freshly propagated liquid culture *C. thermophilum* filaments were fixed with 3 % glutaraldehyde (Sigma, Taufkirchen, Germany) in 0.1 M sodium cacodylate buffer (SCP; pH 7.2) for five hours at room temperature. After fixation the samples were rinsed in SCP and postfixed with 1% osmiumtetroxide (Roth, Karlsruhe, Germany) in SCP for one hour at room temperature. Subsequently the samples were rinsed with water, dehydrated in a graded ethanol series, infiltrated with epoxy resin according to Spurr (1969), polymerized at 70°C for 24 hours and then cut to 70 nm ultra-thin sections with an Ultracut S ultramicrotome (Leica, Germany). After cutting, the sections were applied on copper grids with formvar coating and uranyl acetate and lead citrate was added for post-staining in a specialized EM-Stain device (Leica, Germany). For imaging, a Zeiss EM 900 TEM (Carl Zeiss, Germany) operating at 80 keV was used. All image processing was performed using the Fiji software (Schindelin et al., 2012).

### CFS preparation

To prepare the CFS, carefully grown mycelium^43^ was isolated with the use of a 180 μm-pore size sieve, then washed 3 times with PBS at 3 000 g, 4 min and 4°C. After removing any residual moisture, the pellet was freeze-ground using a liquid N_2_ pre-chilled mortar and stored for later usage at -80°C. Approximately 8 g of the freeze-ground material were lysed in 20 mL of Lysis Buffer (100 mM HEPES pH 7.4, 5mM KCl, 95 mM NaCl, 1 mM MgCl_2_, 1 mM DTT, 5% Glycerol, 0.5 mM EDTA, Pefabloc 2.5 mM, Bestatin 130 μM, 10 μg mL^−1^ DNAse, E-64 40 μM, Aprotinin 0.5 μM, Pepstatin A 60 μM, Leupeptin 1 μM), with 3 repeats of 6.5 mps shaking speed for 25 s, 4°C in a Fastprep cell homogenizer and 3 min of rest in ice after each repeat. A 4 000 g centrifugation step was used to pellet the larger cell debris and a 100 000 g high-speed centrifugation was then performed. The resulting supernatant was filtered through a 100 KDa cutoff centrifugal filter and concentrated to 30 mg·mL^-1^.

The filtered, concentrated to 30 mg·mL^-1^ supernatant, was applied to a Biosep SEC-S4000 size exclusion column that is mounted to an ÄKTA Pure 25M FPLC (Cytiva, USA) system via a 500 μL loop. The column was equilibrated with a filtered, degassed buffer containing 200 mM of CH_3_COO^-^NH4^+^ at pH 7.4 prior to sample application. Fraction volume was set to 250 μL and flow rate to 0.15 mL·min^-1^. Based on acquired MS data (see details below), fraction 6 was selected to be tested for suitability as a succinyl-CoA producing CFS.

In order to produce an equivalent yeast sample for enzymatic activity comparison, the *Saccharomyces cerevisiae* strain from ATCC (American Type Culture Collection PO Box 1549 Manassas, VA 20108 USA; ATCC^®^ 24657TM) was cultivated in YPDG medium at 30°C for 5 h, to a OD_595_ of 2.5 (early exponential phase), then harvested at 3 000 g for 5 min at 4°C and washed with distilled water (resulting pellet ∼7 g). The subsequent protocol until final sample preparation is identical to the *C. thermophilum* preparation described above.

### Oxoglutarate dehydrogenase (OGDHc) activity assays

The OGDH activity assay was adapted from^67^. The enzyme assay was prepared in a reaction volume of 100 μl at 4°C, containing 100 mM NaCl, 30 mM K_2_HPO_4_ (pH 7.5), 2 mM MgCl_2_, 2 mM ThDP, 4 mM α-ketoglutarate, 3 mM NAD+, 0.4 mM CoA, and 4 μl Cell Counting Kit 8 and 2 μl cell lysate containing OGDH. For KM calculations, α-ketoglutarate was titrated from 50 to 5,000 μM, NAD+ from 25 to 5,000 μM, and CoA from 5 to 1,000 μM. The reaction mixture (without α-ketoglutarate) was pre-incubated for 5 min at 37 °C for the substrate *K*_M_ calculations and at 25 °C to 65 °C with 5 °C intervals for temperature-dependent kinetic characterization, and the reaction started by the addition of α-ketoglutarate. Formazan product formation by WST-8 of the Cell counting kit 8 was monitored every minute for 1 hour at 460 nm, and concentration calculated with the Lambert–Beer-Equation and the molar absorption coefficient of WST-8^68^ (ε = 3.07 × 104 M^−1^ cm^−1^). Due to substrate excess inhibition, reaction rates were plotted against substrate concentrations using the double reciprocal Lineweaver-Burk plot^69^, and *K*_M_-values were determined at the abscissa intersection point of the asymptotic linear regression.

### Immunoblotting experiments

For Western Blotting (WB) experiments, in-house casted, freshly prepared, 1 mm thickness gels were used with the following composition: separating phase: 370 mM Tris-HCl pH 8.8, 10% w/v acrylamide (37.5:1), 0.04% w/v APS, 0.002% v/v TEMED, 0.1% w/v sodium dodecyl sulfate (SDS) in ddH_2_O, stacking phase: 125 mM Tris-HCl pH 6.8, 5% w/v acrylamide (37.5:1), 0.04% w/v APS, 0.002% v/v TEMED, 0.1% w/v SDS in ddH_2_O. Samples were previously mixed with a 4x loading dye (250 mM Tris-HCl pH 6.8, 40% v/v glycerol, 20% v/v β-mercaptoethanol, 0.2% w/v bromophenol blue, 8% w/v SDS) and then incubated at 100°C for 5 min. For each native sample, around 400 ng of protein was loaded in each lane and 5 μL of Precision Plus Protein™ All Blue Prestained Protein Standards (Biorad, USA) was loaded in each gel as a marker. Concentration of recombinant control samples was around 8 ng. After loading, gel electrophoresis followed in a 1X electrophoresis buffer, freshly diluted from a 10X stock (144 g Glycine, 30.3 g Tris-base in ddH_2_O) with an applied electrical field of 100 V for 1,5 h. A Trans-Blot^®^ Turbo Transfer System (Biorad, USA) was used to transfer the gel contents to a nitrocellulose membrane for 20 min, with a pre-set, 25 V (1 A) applied field. Blocking was performed for 1 h, under constant stirring in 5% w/v TBST/milk solution and then the membranes were incubated at 4°C for 16 h with the primary antibody (0.2 μg·ml^-1^, 2% w/v TBST/milk). The primary antibody was removed and three washing steps of 2% w/v TBST/milk followed. The membrane was then incubated with the secondary antibody (Goat Anti-Rabbit IgG, Abcam, ab205718, 0.1 μg·ml^-1^, 2% w/v TBST/milk) for 1 h. Three more washing steps of 2% w/v TBST/milk were applied and finally the membranes were screened with a ChemiDoc MP Imaging system (Biorad, USA) and freshly mixed ECL fluorescent mixture and optimal exposure times. All primary antibodies were tailor-made by GenScript (New Jersey, USA) with sequences found in **Supplementary Table 7**.

### Cryo-EM sample preparation and data collection

For structural characterization of the succinyl-CoA producing CFS, a 3.5 μL sample of final protein concentration of 0.3 mg·mL^-1^ was applied on a carbon-coated, holey support film type R2/1 on 200 mesh copper grid (Quantifoil, Germany) that was previously glow discharged under the following conditions: 15 mA, grid negative, 0.4 mbar and 25 s glowing time with a PELCO easiGlow (TED PELLA, USA). The grid was then plunge-frozen with a Vitrobot^®^ Mark IV System (ThermoFisher Scientific, USA) after blotting with Vitrobot^®^ Filter Paper (Grade 595 ash-free filter paper ø55/20 mm). In the chamber, conditions were stabilized at 4°C and 95% humidity, while blotting parameters were set at 0 blot force and 6 s of blotting time. The vitrified grid was clipped and loaded on a Glacios 200 keV Cryo-transmission electron microscope (ThermoFisher Scientific, USA) under cryo and low humidity conditions. Images were acquired with the Falcon 3EC direct electron detector and the EPU software (ThermoFisher Scientific, USA) in linear mode and total electron dose of 30 e-/Å^2^. Before acquisition, the beam was aligned to be parallel and perpendicular to the sample, with a 2.5 μm diameter, while a 100 μm objective aperture restricted the objective angle. Complete acquisition parameters are listed in **Supplementary Table 1**.

### Image processing

All steps of image processing were performed with the cryoSPARC high-performance computing software version 3.3.1^70^. A dataset of 25 803 movies was imported to a dedicated workspace. The in-software patch motion correction (multi) and patch CTF estimation (multi) algorithms were employed to correct for beam induced motion and calculate for the CTF parameters of the micrographs respectively. After dataset curation, 24 300 micrographs were selected for subsequent steps of the image analysis process. Templates were created from EMD-13844^18^ with the create templates job (20 equally-spaced generated templates) and were employed for a picking job with the template picker, resulting in an initial dataset of 3 596 302 particles that was extracted with a box size of 208 pix. and then reference-free 2D classified in 200 classes. From the initial 2D classification, 71,912 particles were selected from 4 classes and subjected to heterogeneous refinement, further refining the final particle set to 52 034 particles after discarding particles that resulted in mal-formed OGDHc E2o core structures. The final particle set was then symmetry expanded with octahedral (O) symmetry and employed for the final core reconstruction with the local refinement (new) job, resulting in a OGDHc E2o core map of 3.35 Å resolution (Gold-standard FSC criterion 0.143) (**Supplementary Figure 2A**). For the reconstruction of the complete OGDH complex, the 71,912 particles that were included in the core reconstruction were re-extracted with a larger box size of 288 pix in order to include signal for the subunits that are located in the periphery of the core. They were then re-classified in 20 2D classes and the classes that showed most prominent peripheral densities were used again as template for a new round of template picking, resulting in an initial particle set of 2 891 518 single particles. This particle set underwent 3 more rounds of 2D classification, always selecting towards class averages that displayed a robust core signal, but in addition to the core also displayed peripheral subunit signal, ending up with a set 52,551 particles that were finally used for a 3D classification job with 10 classes. Class 0, containing 5,178 particles and displaying the most well-resolved peripheral densities, was finally used for homogeneous refinement, resulting in a OGDHc map of 21.04 Å resolution (Gold-standard FSC criterion 0.143) (**Supplementary Figure 2B**) which was then utilized for all structural analysis. All map visualization was performed with the ChimeraX^71^ software package.

### Atomic model building and refinement

For refinement of the E2o core structure, the initial model (PDB ID: 7Q5Q) was fitted into the cryo-EM density using ChimeraX and then refined using iterative manual refinement with Coot^72^ and real-space refinement with Phenix^73^ with standard parameters. Visible density that could be mapped was extended towards the *N-ter* of the E2o, from M195 to E188.

### AI-based model generation and electrostatic surface calculation

A local installation of AlphaFold2-Multimer^74^ was utilized in order to perform predictions of the E2o core vertex trimer that was modelled in the experimentally derived map, the E1o dimer (Uniprot ID: G0RZ09) and E3 dimer (Uniprot ID: G0SB20) in complex with the Uniprot-annotated E2o LD domain (residue numbers 40 to 115, Uniprot ID: G0SAX9). All model validation metrics can be found in the **Supplementary Figure 5A, B**. In order to calculate and visualize all electrostatic surfaces, the APBS electrostatics plugin^75^ was used in PyMol (Schrödinger, USA).

### Energetic calculations and macromolecular docking

Two distinct protocols were applied for reported HADDOCK2.2 calculations: (a) HADDOCK refinement^76^. Here, the refinement protocol was applied to calculate and compare the energetics of interfaces between OGDHc components and its human homologue as well as the Alphadfold-generated interfaces formed by the LD. In this refinement procedure, only the water refinement stage of the HADDOCK protocol was performed that showed to qualitatively correlate calculated energetics with binding affinities for transient protein-protein interactions^40^ skipping the docking step. For this, complexes were solvated in an 8Å shell of TIP3P water. The protocol consisted of the following steps: (1) 40 EM steps with the protein fixed (Powell minimizer) and (2) 2X 40 EM steps with harmonic position restraints on the protein (k=20 kcal·mol^-1^ Å^−2^). For the final water refinement, a gentle simulated annealing protocol using molecular dynamics in Cartesian space is introduced after step (2). It consists of: (1) Heating period: 500 MD steps at 100, 200 and 300K. Position restraints (k=5 kcal·mol^−1^ Å^−2^) are applied on the protein except for the side-chains at the interface. (2) Sampling stage: 1250 MD steps. Weak (k=1 kcal·mol^−1^ Å^−2^) position restraints are applied on the protein except for the backbone and side-chains at the interface; and (3) Cooling stage: 500 MD steps at 300, 200 and 100K. Weak (k=1 kcal·mol^−1^ Å^−2^) position restraints are applied on the protein backbone only except at the interface. A time step of 2fs is used for the integration of the equation of motions and the temperature is maintained constant by weak coupling to a reference temperature bath using the Berendsen thermostat^77^. The calculations were performed with CNS^78^. Non-bonded interactions were calculated with the OPLS force field^79^ using a cutoff of 8.5Å. The electrostatic potential (E_elec_) was calculated by using a shift function while a switching function (between 6.5 and 8.5Å) was used to define the Van der Waals potential (E_vdw_). The structure calculations were performed on the HADDOCK web server at https://alcazar.science.uu.nl/ using the refinement interface. A total of 200 structures was generated for each complex. (b) HADDOCK flexible docking. The HADDOCK docking server was used, utilizing the guru interface. Here, distance restraints were used by applying the derived crosslinking data matching LYS residues of the LD and the E1, and E3, respectively. The distance restraints applied are included in the haddockparam.web files provided with this article. For the docking calculations, the guru interface was utilized by default, but search and scoring space were significantly expanded. This meant that it0 generated structures were increased to N=10 000 and scored structures were increased to N=400. Calculations of frequent amino acid residues involved in the binding of the LD were produced from the formed interfaces from the N=400 final, water-refined docking solutions. An interface residue is considered if it is in proximity of 5 Å from any other residue from the LD. Highly frequent residues reported are the ones that belong to the top quartile (above 25%) in the calculated frequencies per residue and these were plotted with the BoxPlotR online webserver^80^.

### Peripheral subunit fitting

The OGDHc complex map that was generated from the cryo-EM experimental data was used to identify the placement of the peripheral E1o and E3 subunits, in complex with the E2o LD domain that were generated through AlphaFold2-Multimer predictions as follows: the map was displayed in ChimeraX and after fitting the E2o core in the center, it was segmented into a “core” region and an “external” density region with the segment map tool. Then a fit search of 100 fits was performed for both E1o-LD and E3-LD complexes and cross-correlation (CC) values for each fit were listed and separated as fitting in the core or external density region. For statistical significance, the two CC fit groups were tested with single-factor analysis of variance (ANOVA) with the Analysis ToolPak in Microsoft Excel (Microsoft Corporation, USA) and values for the fits can be found in **Supplementary Figure 4A, B** and **Supplementary Table 3**. CC plots were plotted with the BoxPlotR online webserver.

### Linker distance and rotational displacement calculations

*C. thermophilum* E2o linker distance calculations were performed as follows: (a) For the experimentally resolved distance measurements, after all peripheral subunits with bound LD were fit in the OGDH complex map, distance was measured with the “distance” command in ChimeraX, starting from the last resolved *N-ter* residue for each of the 3 E2o subunits of the visually inspected closest E2o core vertex trimer, to the first *C-ter* residues of the modelled LD domain bound to either E1o or E3 dimers on the periphery. All values were then averaged, and individual measurements and standard deviations calculated can be found in **Supplementary Table 3**. Theoretical calculations based on disordered linker length (73 a.a., Uniprot ID: G0SAX9) were then based on derived equations published by Marsh and Forman-Kay^55^, Wilkins *et al*.^81^ and George and Heringa^56^. Additionally, to further derive insights on disordered linker length based on experimentally resolved structures from the PDB, the complete PDB database, as of 1st Jun. 2022, was downloaded and a total of 191144 mmCIF format files, containing more than half a million chains were analyzed. Missing linker regions with their corresponding sequences are identified from the “_pdbx_unobs_or_zero_occ_residues” entries in the mmCIF files and a total of 399 404 missing regions were recorded. For each missing region entry, the left and right observed amino acids are identified using the atomic coordinate data in the mmCIF files. For every linker region, following properties are derived (a) the end-to-end Ca-Ca distance between the two observed residues, measured in Å, (b) the length of the linker region, and (c) the sequence of the linker region. Lengths of these missing regions varied from 1 amino acid up to 3736 amino acids, and a sharp reduction of available PDB entries was observed upon increased length of linker sequence, with the trend visible in **Supplementary Figure 4E**. To derive better collective properties, the files were grouped together by adaptively increasing the bin width of length of amino acids. For 1 to 50 amino acids, the bin width was kept at 1. For 50 onwards, the right bin-edge increases gradually, to 55, 60, 65, 70, 75, 100, 125, 150, 250 and 4 000. From entries falling in every bin, the mean value of the Ca-Ca distances was calculated, and distance at quantiles varying from 0.0 to 1.0 in steps of 0.2. The analyzed data is presented in **Supplementary Table 4**. In **Fig. 6A**, the mean Ca-Ca distance, represented by black dots, was plotted against the length of the linker region. Distances at quantile level (0.0 to 1.0) was plotted as a color shade as annotated in the legend. For reference, a horizontal line (dotted blue) was plotted corresponding to 3.5 A. Similarly, a line (solid blue) corresponding to 7 A times the length of the missing region was plotted. The relation between the mean Ca-Ca distance (y) and the number of amino acids of the missing sequence (x) can be empirically characterized by a model function of the form: *y* = *A*(1 − *e*^−*bx*^) + 3.5, where, constants A and b are observed to have values, 11 and 0.2, respectively. The model function was plotted as a red line. All plots related to PDB distance calculations (**Fig. 6A, Supplementary Figure 4D, E**) were generated with the Pandas package in Python 3.9. The bubble plot containing all theoretical and experimental distance calculations (**Supplementary Figure 4C**) was generated with the ggpubr 0.4.0 package in R.

Rotational displacement calculations of the *H. sapiens* E2o vertex trimer subunits vs. the experimentally resolved *C. thermophilum* E2o vertex trimer subunits (A, B, C) were performed as follows: first, *C. thermophilum* and *H. sapiens* (PDB ID: 6H05) E2o trimers were extracted and aligned upon subunit A in PyMol. Then, the “angle_between_domains” command was used to first calculate the angle between *C. thermophilum* subunit A and B. The same was done for the angle between *C. thermophilum* subunit A and *H. sapiens* subunit B. Then the values were subtracted. The same process was done for the subunit pair A and C, again keeping the rotation axis the same, aligned on subunit A. A visual representation of the measured domains can be seen in **Supplementary Figure 3A**.

### Mass spectrometry protein identification and crosslinking mass spectrometry

Fractions 3 to 9 from the *C. thermophilum* native lysate fractionation described above were pooled into two pools (3-6 and 7-9). From each pool, 40 μL of sample was digested in-solution with trypsin as described previously^82, 83^. To avoid protein precipitation during sample reduction and alkylation, 2 μL of 20% SDS were added. 1 μL of 200 mM DTT in 200 mM HEPES/NaOH pH 8.5 was added to reduce the protein samples, which were then incubated at 56°C for 30 min. After reduction, alkylation followed with the addition of 2 μL of 400 mM chloroacetamide in 200 mM HEPES/NaOH, pH 8.5 and another incubation at 25°C for 30 min. All excess chloroacetamide was quenched by adding 2 μL of 200 mM DTT in HEPES/NaOH, pH 8.5. After reduction and alkylation, the samples were used for single-pot solid-phase-enhanced sample preparation^82, 83^. For this preparation, 5 μL of 10% v/v formic acid and 2 μL of Sera-Mag Beads were added, along with enough acetonitrile (ACN) to achieve a final ACN percentage of 50% v/v, then followed by an 8 min incubation and bead capture on a magnetic rack. The beads were then washed two times with the addition of 200 μL 70% ethanol and one more time with 200 μL of ACN. After resuspension in 10 μL of 0.8 μg of sequencing grade modified trypsin in 10 μL 100 mM HEPES/NaOH, pH 8.5, the beads were incubated overnight at 37°C. The incubation was followed by a reverse phase cleanup step and then analyzed by liquid chromatography coupled to tandem mass spectrometry (LC-MS/MS) with a Q Exactive™ Plus Hybrid Quadrupole-Orbitrap™ Mass Spectrometer (ThermoFisher Scientific, USA). More specifically, an UltiMate™ 3000 RSLCnano System (ThermoFisher Scientific, USA) equipped with a trapping cartridge and an analytical column was used for peptide separation. For solvent A 0.1% v/v formic acid in LC-MS grade water and for solvent B 0.1% v/v formic acid in LC-MS grade CAN were used. All peptides were loaded onto the trapping cartridge with a solvent A set flow of 30 mL·min^-1^ for 3 min and eluted with a 0.3 mL·min^-1^ for 90 min of analysis time, starting with a 2-28% solvent B elution, then increased to 40% B, another 80% B washing step and finally re-equilibration to starting conditions. The LC system was directly coupled to the mass spectrometer using a Nanospray-Flex ion source and a Pico-Tip Emitter 360 μm OD x 20 μm ID; 10 μm tip. The mass spectrometer was operated in positive ion mode with a spray voltage of 2.3 kV and a capillary temperature of 275°C. Full scan MS spectra with a mass range of 350– 1400 m/z were acquired in profile mode using a resolution of 70,000 [maximum fill time of 100 ms or a maximum of 3e6 ions (automatic gain control, AGC)]. Fragmentation was triggered for the top 20 peaks with charge 2 to 4 on the MS scan (data-dependent acquisition) with a 20 s dynamic exclusion window (normalized collision energy was 26). Precursors were isolated with 1.7 m/z and MS/MS spectra were acquired in profile mode with a resolution of 17 500 (maximum fill time of 50 ms or an AGC target of 1e5 ions). For the data analysis, the MS raw data were analyzed by MaxQuant 1.6.1^84^. Chaetomium thermophilum proteomes sequences were downloaded from Uniprot with Proteome ID UP000008066. The MS data were searched against Chaetomium thermophilum proteomes sequences plus common contaminants sequence provided by MaxQuant. The default setting of MaxQuant was used with modification oxidation and acetyl (protein N-term). A false-discovery rate (FDR) cutoff of 1% was used for protein identification, and iBAQ intensity was used for label-free protein quantitation. When calculating iBAQ intensity, the maximum detector peak intensities of the peptide elution profile were used as the peptide intensity. Then, all identified peptide intensities were added and normalized by the total number of identified peptides.

For the preparation of the crosslinking mass spectrometry samples, a titration to identify the optimal crosslinker concentration was first performed (**Supplementary Figure 1C**). Fractions 3 to 9 from the *C. thermophilum* native lysate fractionation described above were pooled to a total volume of 1.4 mL and protein concentration of 0.48 mg·mL^-1^, then split into 7 parts of equal volume in each Eppendorf tube. In the first tube no crosslinking agent was added to be kept as control, while to the rest, 5 μL of 0.16, 0.32, 0.63, 1.25, 2.5 and 5 mM of the crosslinking agent BS_3_ was added respectively. The samples were incubated for 2 h on ice and the crosslinking reaction was then deactivated with the addition of 50 mM NH_4_HCO_3_, incubated again for 30 min on ice. The samples were then transferred in an acetone-compatible tube and 4 times the sample volume of cold (−20°C) acetone was added, the tubes were vortexed, again incubated for 60 min at -20°C and centrifuged for 10 min at 15,000 g. The supernatant was properly removed and then the tubes were left open at RT in order for the acetone to evaporate. To visualize the titration results, each of the 7 tubes’ pellets were resuspended in 1X SDS-PAGE sample loading buffer (diluted from a 4X stock of 250 mM Tris–HCl pH 6.8, 8% w/v sodium dodecyl sulfate (SDS), 0.2% w/v bromophenol blue, 40% v/v glycerol, 20% v/v β-mercaptoethanol) to a final protein concentration of 10/20 μg·mL^-1^. The samples were then boiled for 5 min at 90°C, loaded onto Mini-PROTEAN^®^ Precast Gels (BioRad, USA) and electrophorized for 60 min at 150 V. After the electrophoresis, gels were washed twice in water, stained with Coomassie staining solution and then destained until the background of the gel was fully destained. After visual inspection of gels, an optimal concentration of 1 mM BS_3_ was selected to proceed with sample crosslinking (**Supplementary Figure 1C**). With optimal crosslinker concentration determined, fractions 3 to 9 from the *C. thermophilum* native lysate fractionation described above were pooled again, but this time in two pools, 3-6 and 7-9. In a Protein LoBind Eppendorf tube for each pool, 100 μL of 8M urea was added and then centrifuged overnight in a centrifugal vacuum evaporator at RT. 100 μL from each pool was then transferred to each of the tubes containing urea powder and 100 mM of ammonium bicarbonate (ABC) was added. DTT was then also added to a final concentration of 2.5 mM (from a 100 mM DDT stock dissolved in 100 mM ABC) and incubated for 30 min at RT. After the incubation was over, 2-Iodoacetamide (IAA) was added to a final concentration of 5 mM followed by incubation for 30 min, at RT in the dark. The reaction was quenched by the addition of 2.5 mM DTT to prevent modification of serine in the trypsin active sites. LysC (1μg μL^-1^ stock, in 1:100 ratio, w/w) was subsequently added and samples were again incubated at RT for 4.5 h. An addition of 50 mM ABC to the sample reduced the urea concentration to < 2 M. Trypsin (from 1 μg μL^-1^ stock, in 1:50 ratio, w/w) was introduced to the samples which were then incubated overnight at RT, followed by STop And Go Extraction (STAGE) TIPS desalting procedure, as described in^85^. The final crosslinked peptide samples were subjected to the same LC-MS/MS process and the results were analyzed as described above in the MS protein identification method. All plots, visualizing MS/XL-MS data were created with the xiVIEW online webserver^86^.

### Network analysis and community identification

Identified proteins from the mass spectrometry were mapped in the provided higher-order assemblies described in the Kastritis et al. 2017^34^. These higher-order assemblies were further enriched utilizing homology search per UNIPROT entry against the Protein Data Bank, to also include possible homologous subunits resolved in PDB structures after 2017. For this, Blastp was used (https://blast.ncbi.nlm.nih.gov/Blast.cgi?PAGE=Proteins) by default against the PDB database. Next, networks and communities provided in Kastritis et al. 2017 acted as a basis for further expanding networks reported in this study. This was performed by mapping the proteins to those networks, including them in the STRING^87^ database and expanding the identified interactions into a network with first- and second-shell interactors and communities. Then, *C. thermophilum*-specific local STRING network clusters were retrieved and coverage per string cluster is reported by simple division of identified proteins over all proteins present in the STRING cluster. This was performed from these expanded networks via back-mapping to the MS and XL-MS data. Finally, networks were checked for biological significance utilizing the in-database enrichment detection criterion^88^, showing that all 54 clusters (protein communities) derived here were significantly enriched in biological interactions. Visualizing of the networks was performed with Cytoscape^89^. Recovery of the complete TCA cycle was inferred utilizing KEGG^90^.

### Multiple sequence alignment

Uniprot ID: G0S3G5 (*C. thermophilum* putative holocytochrome C synthase - sequence length: 382), Uniprot ID: Q7S3Z3 (characterized in^31^ as the *N. crassa* Kgd4 protein - sequence length: 130), along with Uniprot ID: Q7S3Z2 (*N. crassa* putative holocytochrome C synthase - sequence length: 317) were downloaded in .fasta format from the Uniprot database and aligned with Clustal Omega^91^ in two separate alignment pairs: G0S3G5 with Q7S3Z3 and G0S3G5 with Q7S3Z2 separately. The .aln files were downloaded and visualized with Jalview^92^. Alignment results are displayed in **Supplementary Figure 5C**.

## Supporting information

List of All PDB-IDs analyzed in the paper

Network Visualization Cytoscape File

## Acknowledgements

We would like to thank all the Kastritis lab members for fruitful discussions.

## Author contributions

F.L.K. carried out the experimental work, assisted by F.J.O. in mass spectrometry and T.K.T in kinetic assays. G.H and M.F performed cell imaging. F.H. acquired the micrographs. I.S. calculated the cryo-EM reconstructions and performed computational modelling. C.T. performed computational modeling. P.L.K., I.H. and J.R. secured funding. P.L.K. conceived the project. I.S. and P.L.K wrote the manuscript with contributions from all authors.

## Data availability

Source data for all figures is provided with this paper in Supplementary Tables. All MS data, cryo-EM data, maps and models will be deposited and will be publicly available. Data essential for reviewers are provided under: https://cloud.uni-halle.de/s/4iDLknhVRK3OxMB with a README file. Folders and files within the zip file include:

- 01_experimental_models
- 02_PDB_validation_e2o_core_map-model_validation_tentative.pdf
- 03_alphafold_models
- 04_HADDOCK_parameters_docking
- 05_Networks_communities.cys
- 06_PDB_files_used_for_Fig6A

## Figure Legends

**Supplementary Figure 1:**
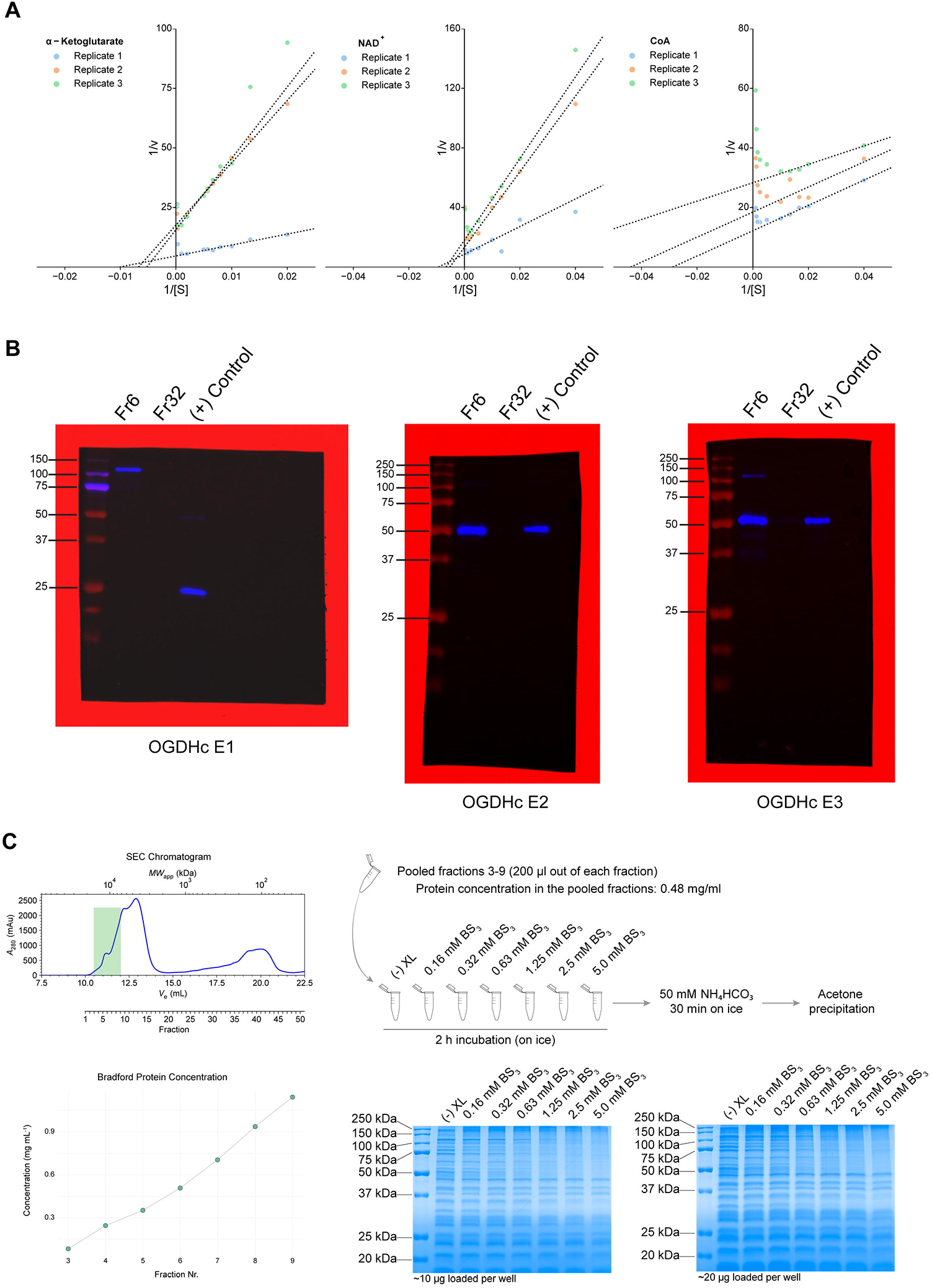
Lineweaver-Burk plots, uncropped WB gels and crosslinker benchmarking. **(A)** Lineweaver-Burk plots for each of the substrates that are involved in the OGDHc reaction, showing the values for the 3 biological replicates for each. **(B)** Un-cropped WB gels for each of the OGDHc components. **(C)** Benchmarking process of the cross-linker concentration. On the top left, the chromatogram of the SEC fractionation of the native *C. thermophilum* lysate that was performed is visible, with the fractions that were pooled for the benchmarking annotated with a green box. Below, protein concentrations for each of the fractions 3-9 are shown. On the right, a schematic representation of the crosslinking process is visible, with the gels that were used to validate the crosslinking process seen on the bottom right.

**Supplementary Figure 2:**
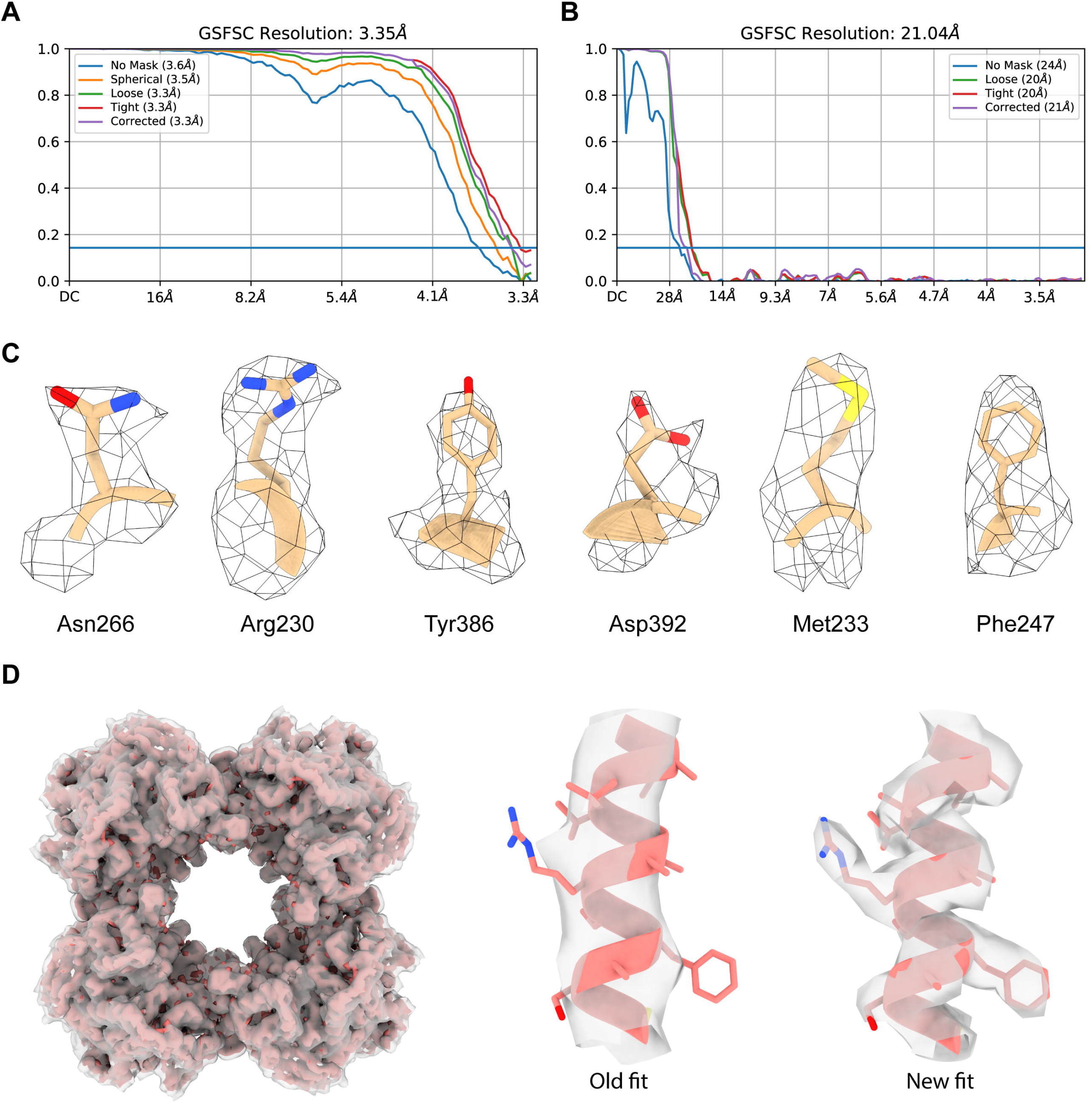
FSC plots and side-chain density resolution examples. **(A)** Gold-standard (0.143) FSC plot for the E2o core map. **(B)** Gold-standard (0.143) FSC plot for the OGDHc map. **(C)** Various examples taken from the cryo-EM resolved fitted model, showing confident placement of side-chains in their corresponding densities. **(D)** Comparison between the CFS-derived and the previously published OGDHc E2o core map (EMD-13844). On the left, the CFS-derived E2o core map (salmon) is fit into the previously derived E2o core map (EMD-13844, gray). On the right, the marked improvement in side-chain density coverage between the old and new maps can be observed.

**Supplementary Figure 3:**
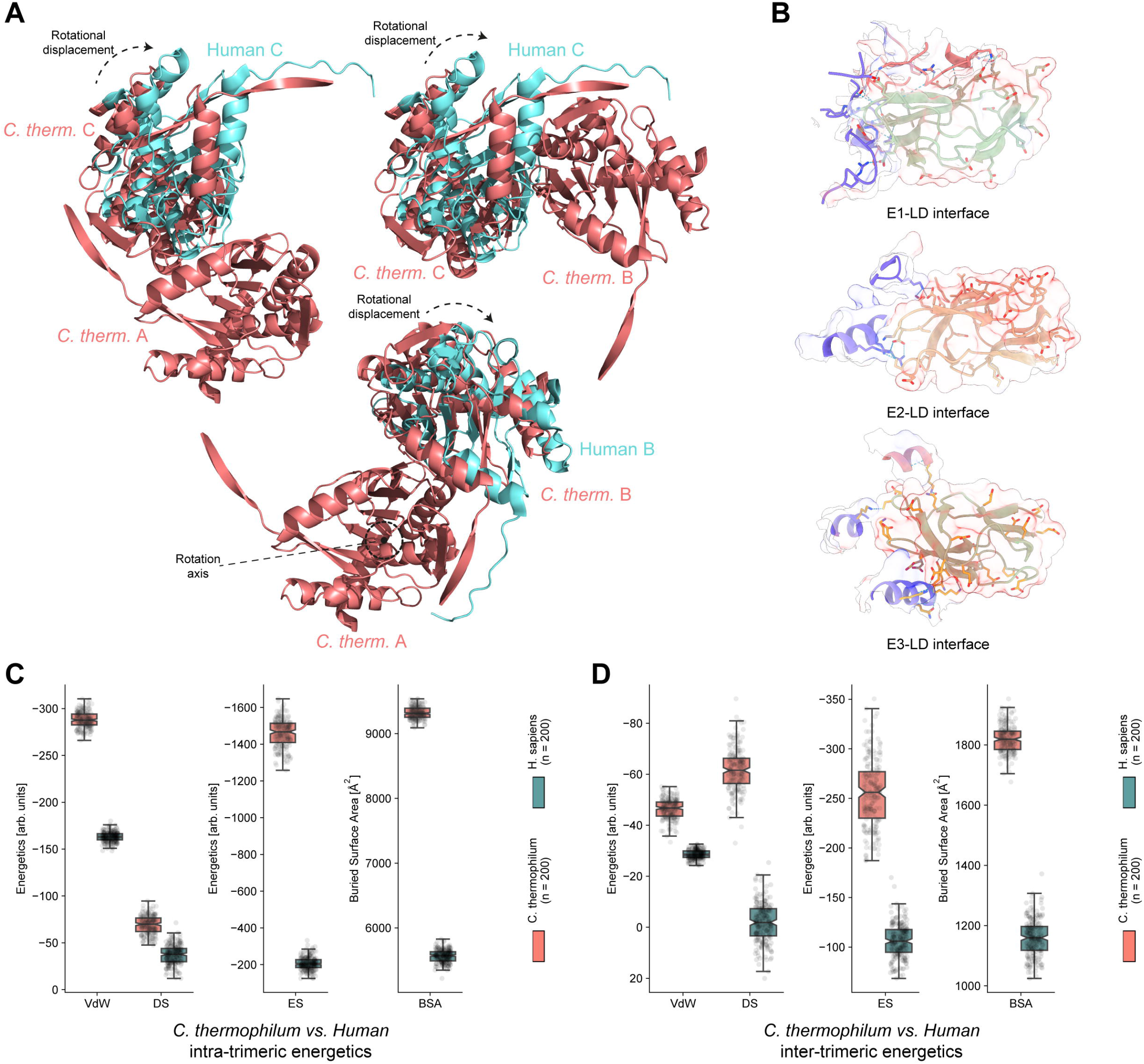
Rotational displacement calculations, interfaces of LD with all OGDHc components and comparative energetics between *C. thermophilum* E2o interfaces and its mesophilic human counterpart. **(A)** Models representing how the rotational displacement calculations were performed, between the E2o core vertex trimer of *C. thermophilum* and Human. **(B)** Residues participating in the interface between the LD and E1o, E2o and E3 respectively. The high number of negatively charged side-chains on the LD surface is visible, along with the hydrogen-bond network that helps stabilize the highly electrostatically-driven interaction. **(C)** *C. thermophilum* and Human E2o core intra-trimeric energetics display stronger forces that contribute to the *C. thermophilum* E2o core compaction. **(D)** *C. thermophilum* and Human E2o core inter-trimeric energetics display stronger forces that contribute to the *C. thermophilum* E2o core compaction.

**Supplementary Figure 4:**
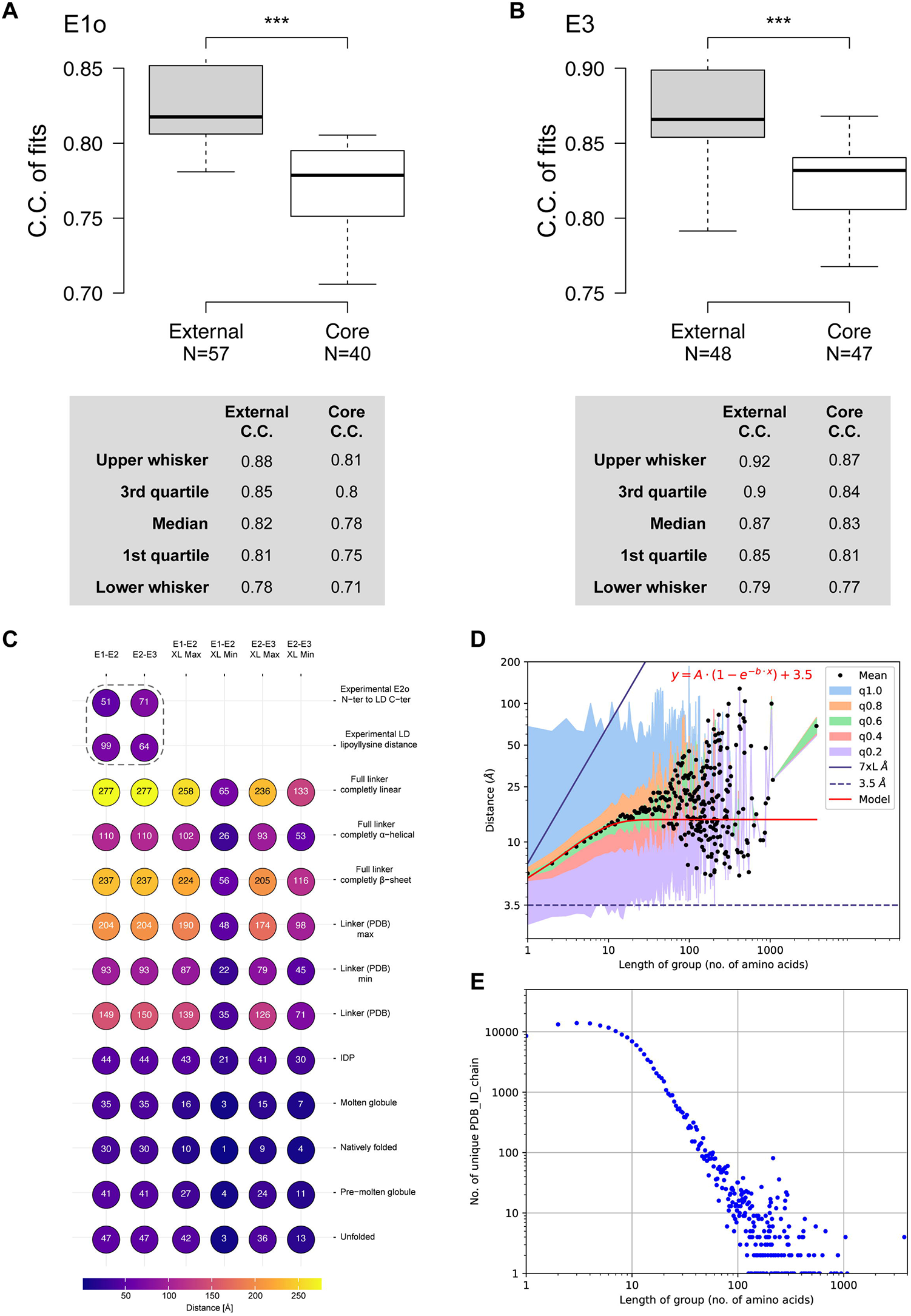
Cross-correlation of fits, experimental and theoretical linker E2o distances, PDB linker distances. **(A)** Boxplot of cross-correlation values of E1o fits in the OGDHc map. The E1o dimer fits to the external map densities with statistical significance compared to the internal densities. **(B)** Boxplot of cross-correlation values of E3 fits in the OGDHc map. The E3 dimer fits to the external map densities with statistical significance compared to the internal densities. **(C)** Bubble plot of all experimental and theoretical E2o linker distance values. XL-min and XL-max annotate the distance from the first *N-ter* residue of the E2o that is resolved in the core map to the closest and farthest cross-linked lysine to a peripheral subunit respectively. **(D)** Graph representing the mean distance values of all unresolved amino-acid sequences belonging to all protein structures deposited in the PDB. Black dots represent the mean value of a length of amino-acids group, with different colors the standard deviation after the integration of each quartile of total data, as denoted in the plot legend. The red line represents a fitted model that describes the relationship between the distance and the length of amino-acid groups. The blue line represents the 2x theoretical distance of an amino-acid sequence, while the dashed blue line represents the theoretical lower limit of any amino-acid sequence. **(E)** Plot representing the trend between the number of PDB structures incorporating a certain sequence length of unresolved amino-acids.

**Supplementary Figure 5:**
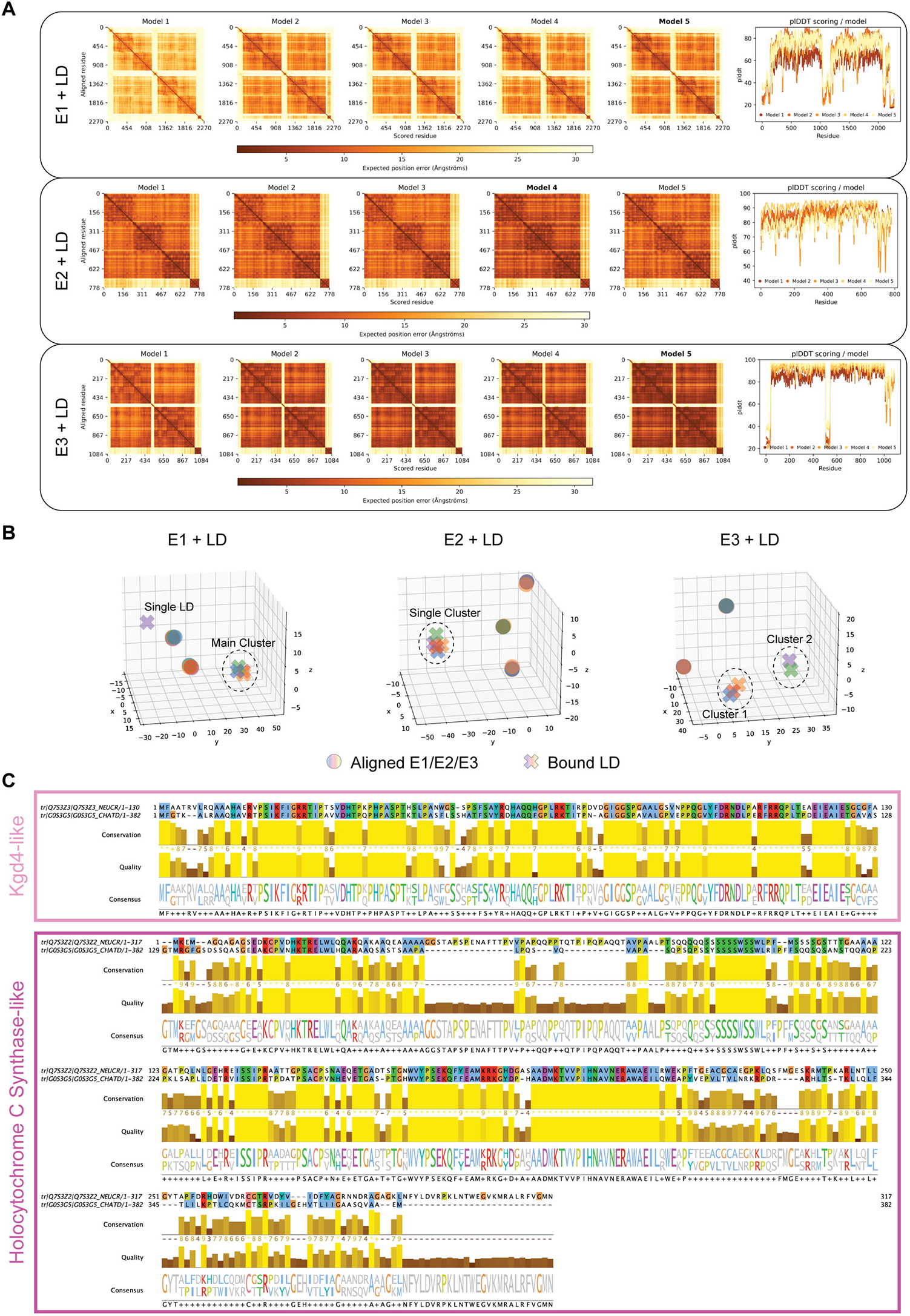
AlphaFold model validation and E3BPo sequence alignments. **(A)** Positional Alignment Error plots and plDDT scoring for the top 5 models of each AlphaFold-multimer prediction of E1o, E2o and E3 in complex with the E2o LD domain. **(B)** Spatial plots showing the position of the LD predicted for each solution returned by AlphaFold-multimer. A single cluster is observed for the predicted interaction between E1o-LD and E2o-LD, whereas 2 clusters can be seen for the E3-LD interaction. **(C)** Sequence alignment of *N. crassa* KGD4 and HCCS to the *C. thermophilum* putative HCCS-annotated protein sequence. The *N. crassa* KGD4 protein sequence aligns with high confidence to the first 129 residues of the *C. thermophilum* sequence, while the *N. crassa* HCCS aligns to the rest 254 residues, strongly indicating that the *C. thermophilum* protein sequence is wrongly annotated and is in reality two separate proteins.

**Supplementary Figure 6:**
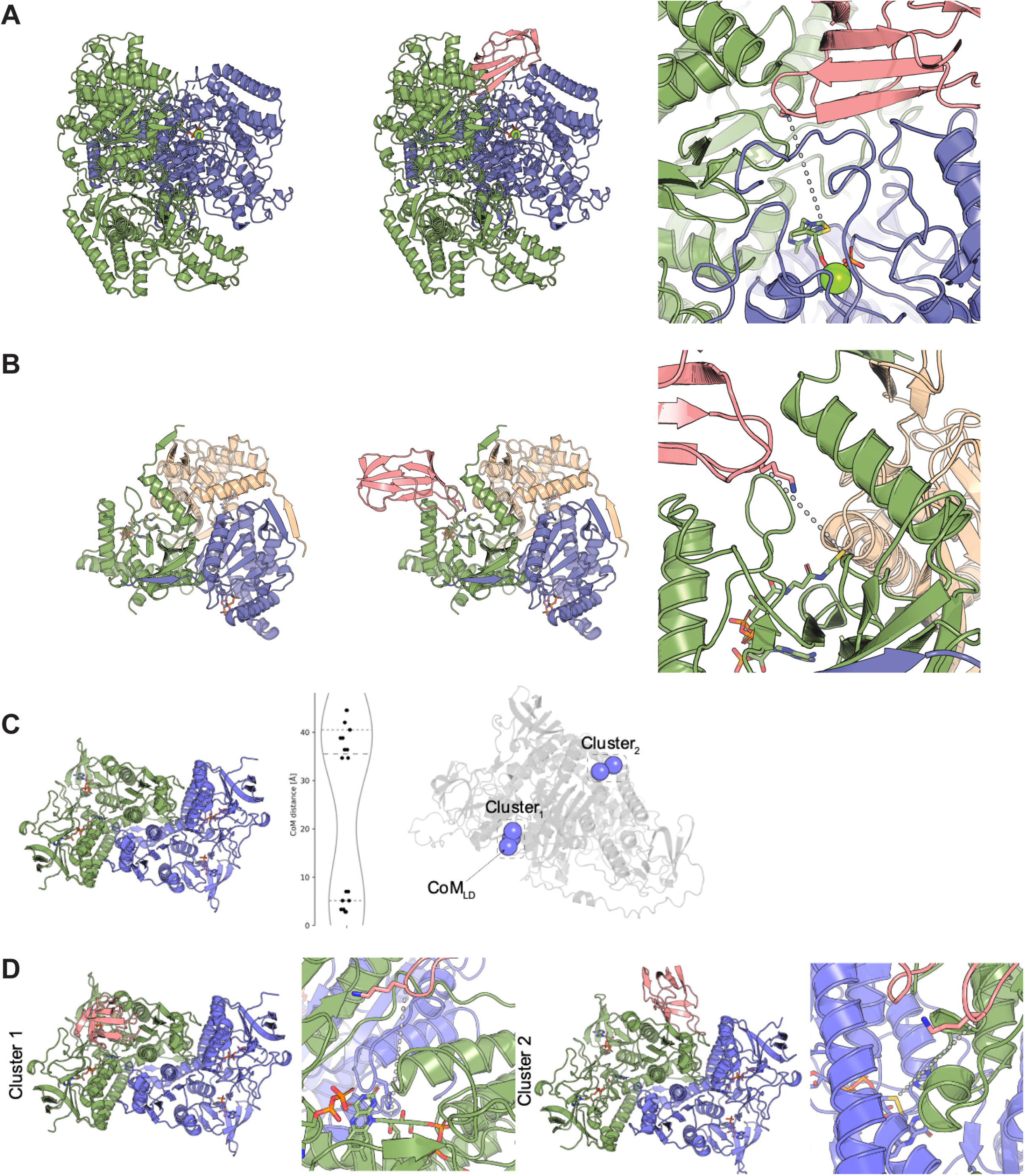
AlphaFold predicted models of the OGDHc subunits in complex with the E2o LD domain. **(A)** AlphaFold predicts the LD as bound in the dimeric interface. The lipoylated lysine is in close distance (14.4 Å) to the C2 atom of the ThDP, which is succinylated during decarboxylation of α-ketoglutarate in the first step of the reaction of OGHDc. The binding cavity of the generated AlphaFold model is not sterically blocked, indicating a plausible docking solution. **(B)** In each dimeric interaction interface in these trimeric E2o building blocks, a CoA binding site is present. Clustering of all AF solutions showed all LDs within a distance range of 10 Å, indicating a single prediction solution. The LD domain is bound to a monomeric E2, meaning that theoretically 24 LD domains could be bound by the OGHDc core simultaneously. The localization of the lipoyl-lysine is close to the CoA binding site, with a distance between the Cα atom of the lysine and the thiol group of CoA, where the succinate from the lipoate is transferred to, of 13.7 Å. The binding cavity is accessible, indicating, again, a plausible docking solution. **(C)** In the predicted structures of E3 with bound LD, there are two clear cluster (n = 3 and n = 2), with clear differences in the localization of the LD. **(D)** Inspecting the two different clusters of the bound LD, in the first cluster, the LD is located at a monomer near the NAD+ binding site, whereas in the second cluster, the LD is bound in the dimeric interface. Mapping the reaction path, in cluster 1, the reaction path is blocked by the FAD, whereas in the second cluster, the lipoylated lysine are in reasonable distance to the disulfide bond in the active side, with a distance of 22.6 Å.

## Tables and table legends

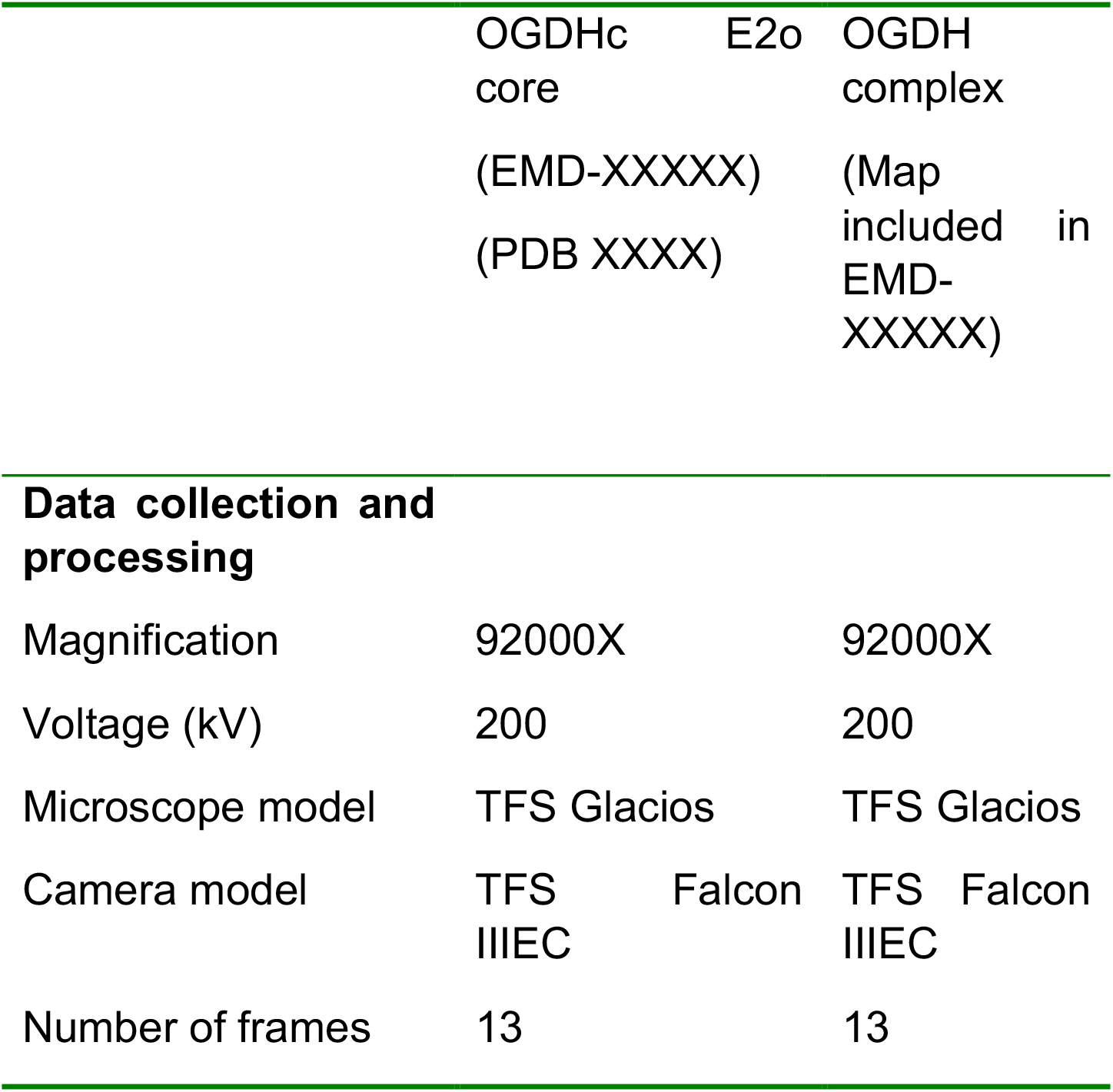
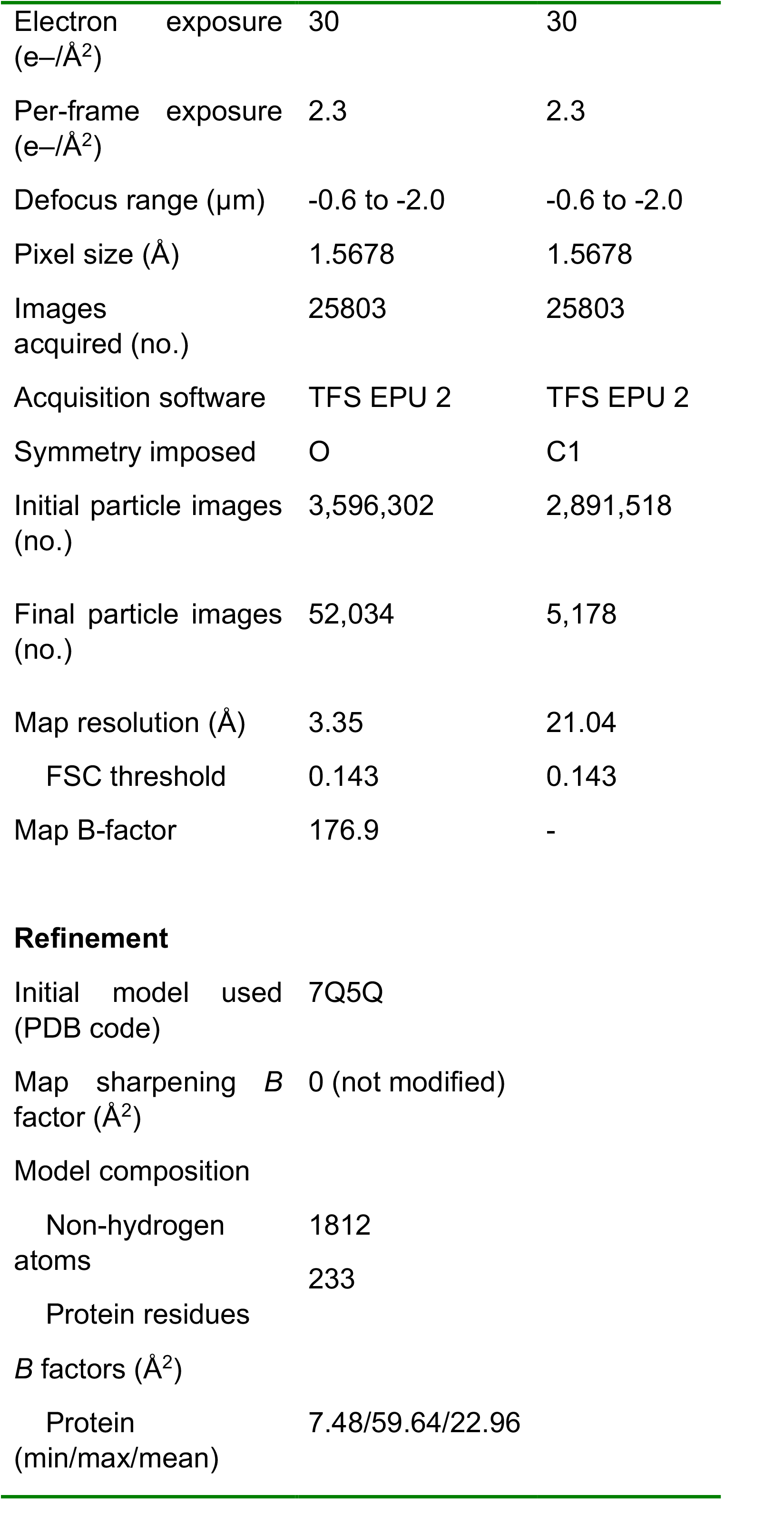
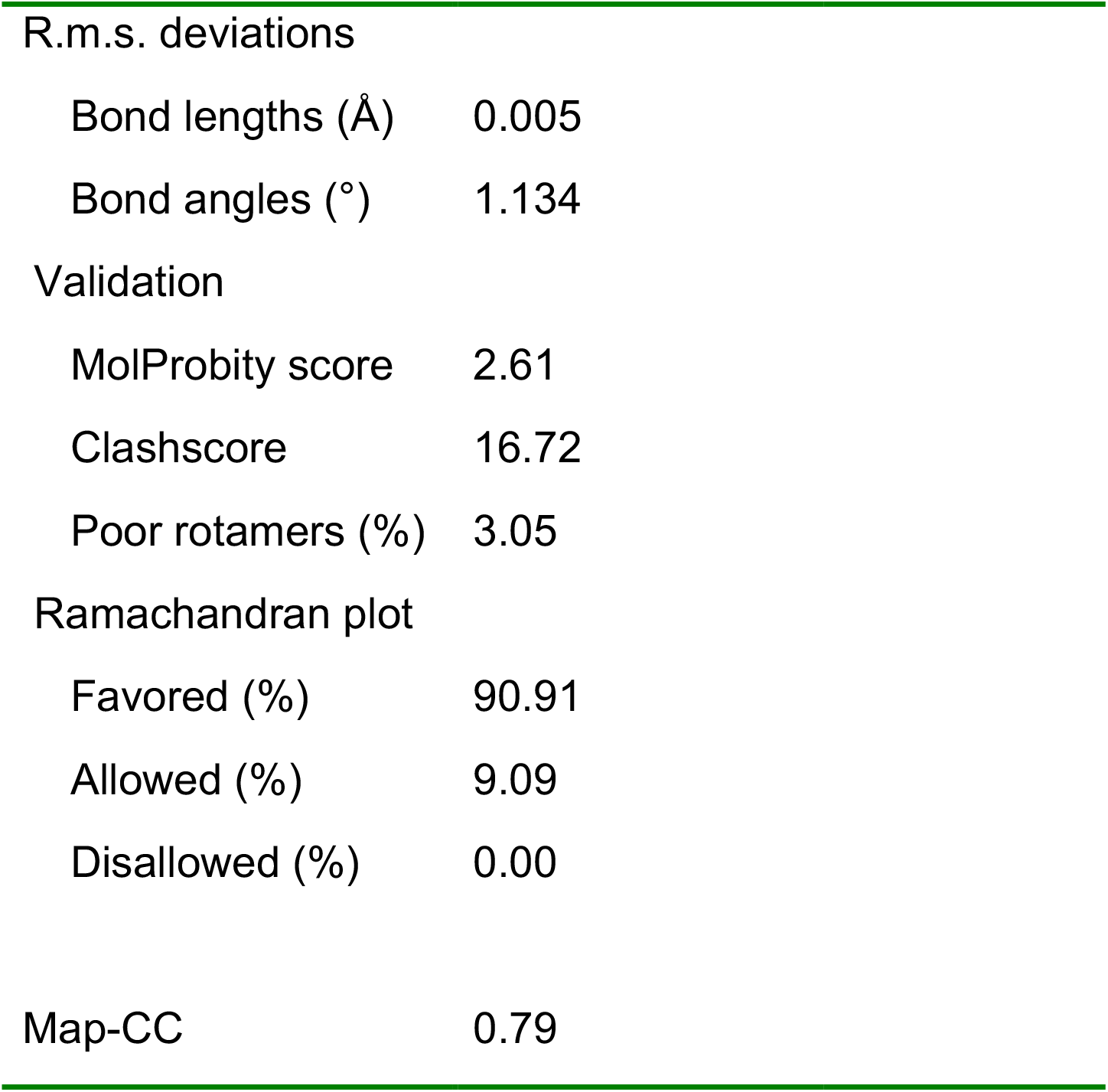

*Supplementary Table 1*: **OGDHc E2o core and complex acquisition and reconstruction parameters**.

*Supplementary Table 2*: **All values related to XL-MS, MS identification, in-fraction community annotation and stoichiometric calculations**.

*Supplementary Table 3*: **All values and statistics related to OGDHc peripheral subunit fits and distance calculations**.

*Supplementary Table 4*: **All values related to plots of unresolved amino-acid sequences in all PDB entries**.

*Supplementary Table 5*: **All values reported in the HADDOCK scoring plots**.

*Supplementary Table 6*: **All values plotted for the kinetic characterization of the OGDHc component reactions**.

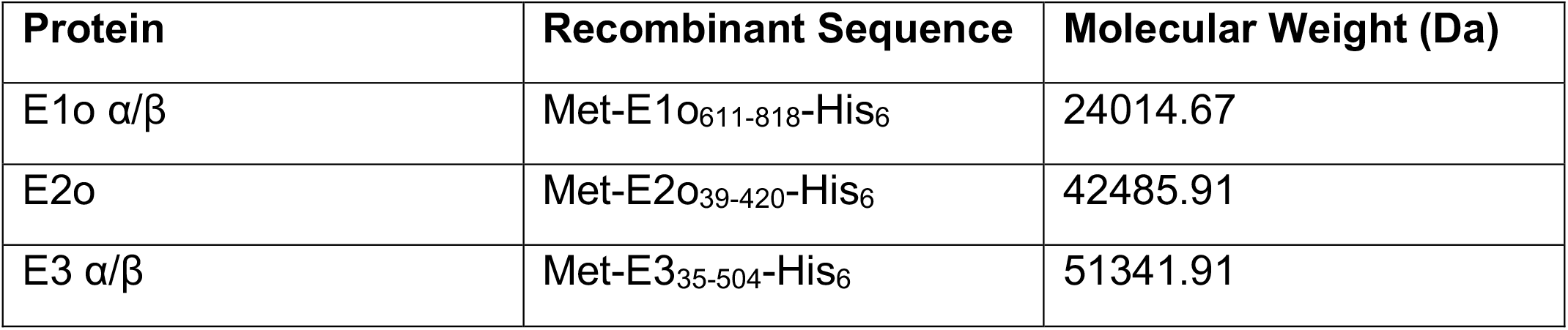

| Protein | Recombinant Sequence | Molecular Weight (Da) |
| --- | --- | --- |
| E1o $\alpha/\beta$ | Met-E1o<sub>611-818</sub>-His<sub>6</sub> | 24014.67 |
| E2o | Met-E2o<sub>39-420</sub>-His<sub>6</sub> | 42485.91 |
| E3 $\alpha/\beta$ | Met-E3<sub>35-504</sub>-His<sub>6</sub> | 51341.91 |

*Supplementary Table 7*: **Sequences of OGDHc components used to produce corresponding antibodies**.

*Supplementary Table 8*: **All residue frequencies of residues participating in the interface between E1o, E3 and the E2o LD domain**.

